# Structural analysis unravels the functional promiscuity of Quinolone synthase-mediated polyketide biosynthesis in *Aegle marmelos* Correa

**DOI:** 10.1101/2022.08.26.505429

**Authors:** Mallika Vijayanathan, KV Abhinav, Debashree Bandyopadhyay, Kozhinjampara R Mahendran, Abdoallah Sharaf, M Radhakrishna Pillai, EV Soniya

## Abstract

Quinolone synthase from *Aegle marmelos* (AmQNS) is a type III polyketide synthase that yields therapeutically effective quinolone and acridone compounds. Based on the high-resolution protein structure of AmQNS, this study provided a mechanistic explanation of the structure to synthetic selectivity. Additionally, it displays the comparatively wide active site entry that allows the catalytic pocket to accommodate bulky substrates, which affects the enzyme catalysis. We also develop a model framework for comprehending the structural constraints on ketide insertion, and postulate that AmQNS synthetic diversity is owing to its steric and electrostatic selectivity, which allows it to bind to a variety of core substrates. We further establish that AmQNS is structurally biased toward quinolone synthesis and only synthesizes acridone when malonyl-CoA concentrations are significantly high. In a nutshell, we anticipate that addressing the structural and molecular underpinnings of AmQNS–substrate interaction in terms of its high selectivity and specificity can aid in the development of numerous novel compounds. Besides, the approaches can also be expanded to other potential enzymes, which will help the pharmaceutical sector by expanding the pool of potential medication leads.

## Introduction

Polyketides (PKs) are chemically diverse natural products with immense pharmaceutical properties. PKs and their possible derivatives could be used as attractive starting points for the development of new bioactive molecules with clinical applications^1,2^. Polyketide synthases (PKS) are multifunctional enzymes that synthesize PKs in plants, fungi, and bacteria^3^. Due to their remarkable characteristic features like i) wide substrate affinity, ii) alternating condensation steps, and iii) formation of diverge cyclic intermediates^4^, the PKS machinery is a good target for producing architecturally diverge natural products by protein engineering and combinatorial biosynthesis. Based on the protein architecture and the mechanism of action, there are three types of PKS, namely type I, II and III. Typically, a polyketide is produced by consecutive addition of ‘malonate building blocks’ to a starter substrate (acyl thioester), catalysed by type III PKS^5,6^. When compared to type I and type II enzymes, type III PKSs are homodimers and comparatively smaller in size. Each functional unit of type III enzyme contains two ketosynthase (KS) domains (∼ 40–45 kDa, ∼350-390 amino acids per monomeric unit)^7–9^. Type III PKSs are further categorized into two subtypes, viz, the i) chalcone-forming (chalcone synthase (CHS)) and ii) non-chalcone-forming (non-CHS), based on the reaction they catalyze^10,11^.

Quinolone synthase (AmQNS)^12^ from the Indian bael tree (*Aegle marmelos* (L.) Correa. or *Crateva marmelos*) belongs to the non-CHS group of type III PKS. *N-*methyl anthraniloyl-CoA is the natural substrate for AmQNS, and the major products are quinolones and acridones. These anthranilic acid-derived quinolone alkaloids (quinine, chloroquine, etc.) have previously been found to have antibacterial, anticancer, and antiviral properties,^13,14^ and could be used as potential pharmacological leads. In general, AmQNS yields diketide 4-hydroxy 1-methyl 2-quinolone (89%) through a one-step condensation reaction between *N*-methyl anthraniloyl-CoA and malonyl-CoA. Acridone (11%) is synthesized in a three-step condensation process that begins with the same substrate and utilizes the same enzyme^12^. When P-coumaroyl CoA is employed as the starting substrate, AmQNS can also produce benzalacetone. The structure and function of two AmQNS-analogue type-III PKSs that use *N*-methyl anthraniloyl-CoA as the starter substrate (*Citrus microcarpa* acridone synthase (CmACS) and quinolone synthase (CmQNS)) have been well characterized^15^. Despite the considerable sequence and structural similarities between AmQNS, CmACS, and CmQNS, their product formation patterns, and catalytic efficiencies are significantly different^12^. i.e., even a minor amino acid substitution can have a significant effect on enzymatic activity. It is remarkable to note that AmQNS and its nearest homolog CmACS both have distinct amino acid variations that favour interaction with the bulky N-methyl anthraniloyl CoA, while hindering the binding of small substrate CoAs. As a result, designing these enzymes to have this mechanical behaviour could improve their biocatalytic characteristics. AmQNS structural and functional features, as well as its reaction mechanism, must be understood in order to optimize its biosynthetic potential for metabolic engineering reprogramming to accelerate natural product discovery.

Here we provide the high-resolution crystal structures of AmQNS in native and substrate-bound forms, as well as the structural and molecular underpinnings for its synthetic selectivity. We used molecular simulations to identify rate-limiting reaction steps leading to quinolone and acridone structure, as well as quantum chemical transition state calculations to compare the relative kinetic barriers and thermodynamic enthalpies of substrates, clearly demonstrating that AmQNS structurally favours quinolone production. Ultimately, these structural findings, together with its simulation-based reaction studies, uncover the mechanistic behaviour of AMQNS and will eventually assist to engineer and repurpose the enzymatic reaction to expand the natural product reservoir for bioprospecting and drug discovery in the future.

## Results and Discussion

### Sequence conservation, evolutionary positioning, and substrate selectivity

In this study, during the homologue search, out of the 283 studied species, type III PKSs were identified in 112 (39.6%) species, and in agreement with previous studies, they are well-conserved in the green lineage (Viridiplantae) and Opisthokonta (Fungi)^3,16^(**Supplementary Table S1C**). Type III PKS were identified in representatives of Rhizaria, Alveolata, and Stramenopila lineages, and its presence on some marine microalgae was also reported previously^17^. It is noteworthy that type III PKS homologs have only been identified in the biflagellated, unicellular, free-living diplonemid *Diplonema papillatum*, among the members of the Discoba - a lineage currently placed proximal to the root of eukaryotes. Besides, we could not identify type III PKS in any of the studied species of Metamonada, Amoebozoa, Glaucocystophyceae, Rhodophyta, Rhodelphea, Rhodelphea, and Haptophyta (**Supplementary Table S1C**). To explore the eukaryotic and prokaryotic type III PKS evolutionary relationship, we traced back to the enzymes in the prokaryotes. Type III PKS were identified in 36 out of the 136 bacterial species studied (26.5%), representing 26 different taxonomic groups (**Supplementary Table S1D**). We searched type III PKS in Archaea and the first time in the Archaeal-Asgard group. As per the previous report^3^, we were unable to identify type III PKS in all archaeal groups but could get homologs in one Asgard species (*Candidatus Thorarchaeota archaeon*) (**Supplementary Table S1D, Supplementary Figure S1**). The Asgard (or Asgardarchaeota) group is a separate domain of life representing the closest prokaryotic relatives of eukaryotes^18,19^. These finding emphasizes the Asgardarchaeota group might be the type III PKS enzyme’s emergence point in the tree of life. In addition to the aforementioned categories, type III PKS homologs from all Rutaceaen species were included in our evolutionary analysis (**Supplementary Table S1B**).

The evolutionary positioning of AmQNS is depicted in the phylogram of all the available homologous proteins (**Figure 1**). AmQNS, like other type III PKS from members of the Viridiplantae, is well conserved and grouped together with other members of the Rutacean family, according to previous evolutionary studies^20,21^. We detected four interesting horizontal gene transfer (HGT) events (**Figure 1**), two of them are within the bacterial domain and close to the root of the tree. First one, between most of the stramenopiles and the cyanobacterial *Rivularia sp*. (WP_015119976.1) homolog. These results show that this class of enzyme has a cyanobacterial origin. The second HGT was between all the alveolates, except *Durinskia baltica* and Chlamydiae homologs. A third HGT was again between Chlamydiae homolog (1444712.BN1013_00142) and the chromerid *Vitrella brassicaformis* homologs; forming a sister group with alveolate *Durinskia baltica*, all fungi, Rhizaria and Discoba homologs and some homologs of stramenopiles (**Figure 1**). A recent report highlighted the contribution of Chlamydiae on the evolution of eukaryotes^22^. Interestingly, the last HGT was in form that all the Planctomycetes and cyanobacterial *Synechococcus sp*. Our analysis shows that this enzyme has a bacterial origin and indicates early origin or even its presence in the Last Eukaryote Common Ancestor (LECA).

**Figure 1.**
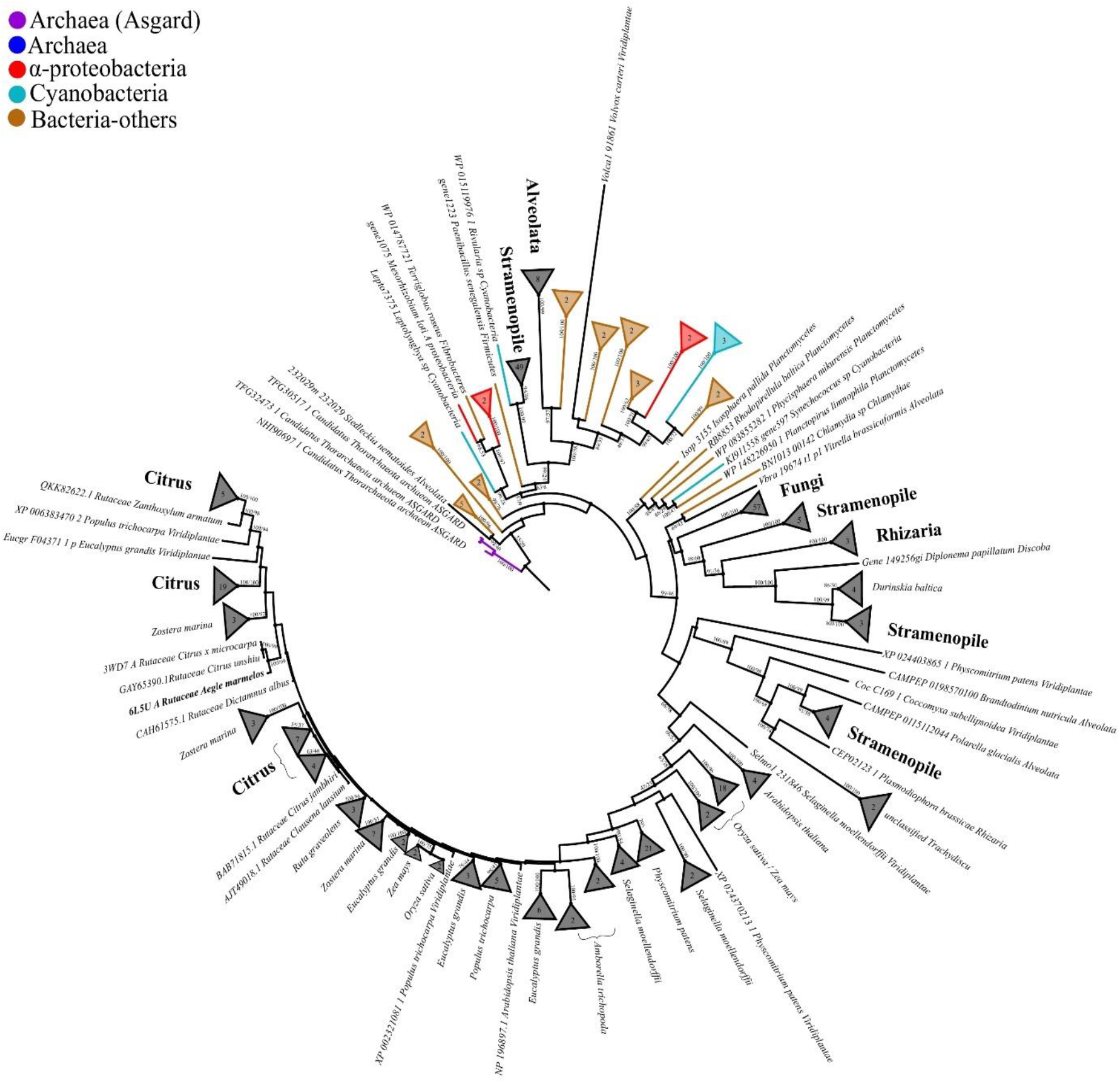
Maximum likelihood (ML) phylogenetic tree of type III PKS protein homologs, showing the potential evolutionary relationships between the identified prokaryotic and different eukaryotic homologs. Phylogenetic position of AmQNS (6L5U) is highlighted in bold. The tree was rooted using the Asgard homologs, to identify the evolutionary direction of the proteins. The maximum likelihood branch support values are given in % (IQ-TREE/RAxML-NG).

Regarding the conservation pattern among type III PKSs, even though structure-based sequence alignment of AmQNS with its adjacent homologs revealed high-level sequence conservation and high functional conservancy (**Figure 2**), minor amino acid differences, particularly in the CoA binding/substrate binding/cyclization pocket area, have a significant impact on substrate specificity, i.e., will change the product formation profile. Because small changes in the three-dimensional structure might occasionally impact substrate selectivity, amino acid moieties in the above-mentioned locations can govern the biological reactions (i.e., protein-ligand interactions).

**Figure 2:**
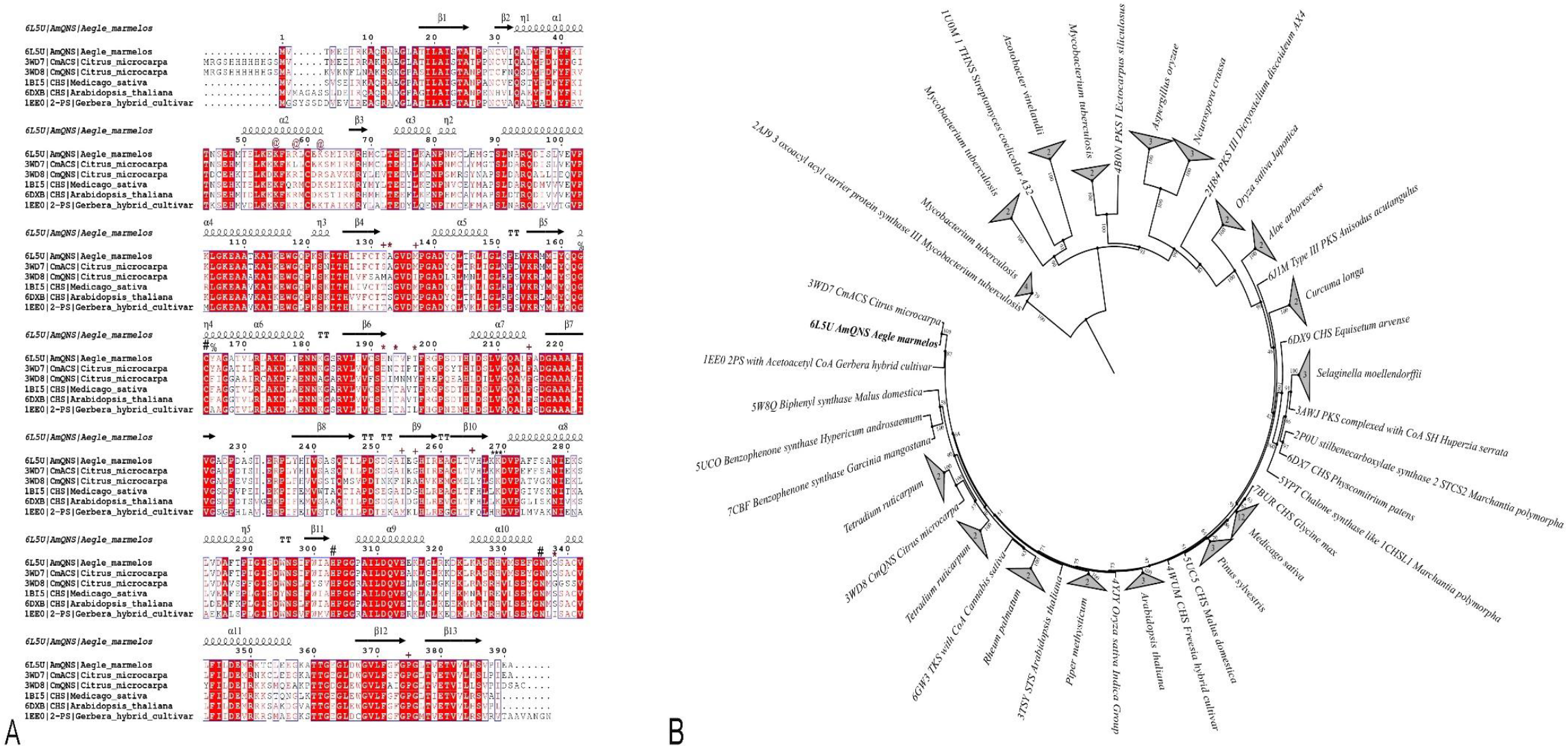
Sequence-structure alignment between different homologs and its evolutionary positions. **A**)Alignment was prepared using ClustalW ^23^ and ESPript 3.0 ^24^. The PDB IDs are used to represent the sequences. The conservation level is shown by a color gradient (white-poor conservation, red-high conservation). Significant residues are highlighted (**#**- Catalytic residues; *- Residues in the substrate-binding pocket; **+**- Residues in the cyclization pocket), @- Residues in the CoA-binding tunnel; %- Residues adjacent to catalytic C164; *- β-turn region of AmQNS **B**) ML phylogenetic tree showing the evolutionary relationship among the structural homologs in RCSB PDB.

One of the most essential aspects of enzymes that determines its unique reaction is the specific interaction between proteins and ligands (such as substrates or cofactors). Radio-TLC experiments have previously shown that AmQNS could accept multiple starter CoAs as potential substrates^12^. Moreover, *in-silico* study also indicates that non-physiological substrates could be employed as potential AmQNS ligands^20^. The binding mechanism of several acyl-CoA substrates (small aliphatic to bulky aromatic) with AmQNS was further authenticated using Surface Plasmon Resonance (SPR) based assays, which enable for real-time monitoring of kinetic parameters^25^. The high affinity and reasonable interaction between the AmQNS and small molecule ligands are indicated by the K_D_ values, which varied from high nanomolar to low millimolar (2 nM-1 mM). AmQNS demonstrated a high affinity for *N*-methylanthraniloyl-CoA, feruloyl-CoA, and hexanoyl-CoA (with KD of 2.04 nM, 9.83 nM, and 7.30 nM, respectively), and it is worth noting that AmQNS prefers bulkier substrates than short acyl-CoAs (**Figure 3, Supplementary Figure S2**). These affinity parameters complement the previously reported interaction studies using a thin-layer chromatography (TLC)^12^, and when comparing the steady state kinetic parameters for AmQNS with different starter substrate CoA’s, it’s notable that Km values are higher than KD for the majority of the substrates (for *N*-methylanthraniloyl CoA-2.93 μM; p-coumaroyl CoA- 3.62 μM; Feruloyl CoA- 9.14 μM). This suggests that catalysis is more rapid than dissociation. These findings imply the prospect of utilizing various substrates to create novel chemical scaffolds, and the enzyme can be further engineered to accommodate various substrates to boost the catalytic versatility.

**Figure 3.**
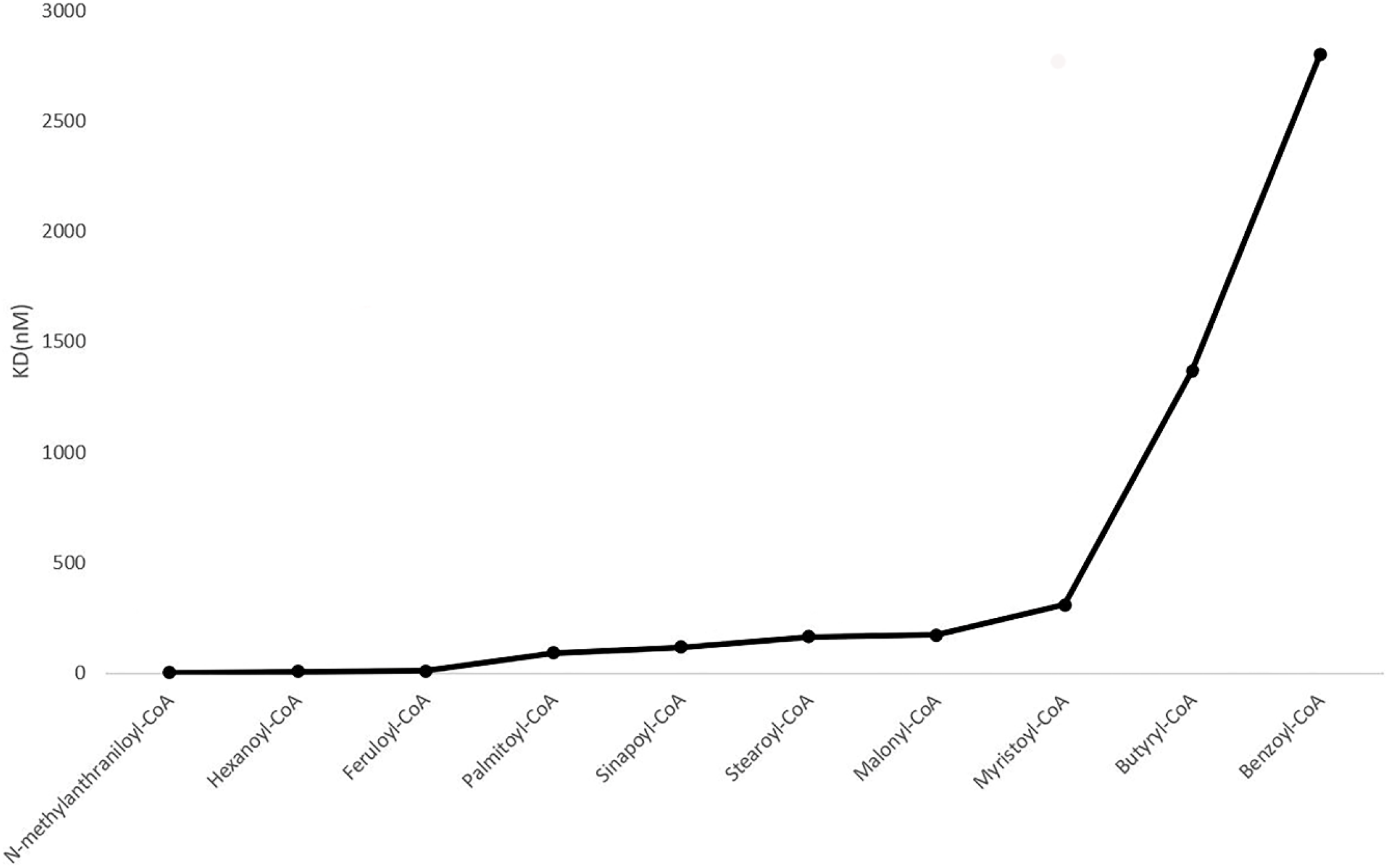
AmQNS interactions with various substrate CoAs. The lower the KD value, the better the interaction.

### Molecular structure of AmQNS in the native and substrate-bound states

The high resolution AmQNS crystals provided very precise protein models with well-defined electron density maps and the structural coordinates have been deposited in the RCSB Protein Data Bank^26^. The data collection and refinement statistics are given in **table 1**.

**Table 1.** Crystallization, data collection and refinement statistics of different crystal forms obtained for native, substrate and CoASH bound AmQNS.

|  | **Native (6L5U)** | **CoASH (6L7J)** | **Substrate bound (7CCT)** |
| --- | --- | --- | --- |
| **Condition** | 0.1 HEPES (pH-7.5), 1.4 M sodium citrate tribasic dihydrate | 0.1 HEPES (pH-7.5), 1.4 M sodium citrate tribasic dihydrate | 0.1 HEPES (pH-7.5), 1.4 M sodium citrate tribasic dihydrate +2mM *N*-methylantraniloyl-CoA (soaked) |
| **Additive** | 0.1M magnesium chloride hexahydrate | 0.1M BaCl<sub>2</sub> | 0.1M Cadmium chloride hydrate |
| **Protein concentration (mg/ml)** | 10 | 10 | 10 |
| **Cryoprotectant** | 20% glycerol | 20% glycerol | - |
| **Space group** | H32 | H32 | H32 |
| **Number of molecules/ASU** | 1 monomer | 1 monomer | 1 monomer |
| **Unit cell dimensions** |  |  |  |
| a (in Å) | 149.84 | 150.86 | 148.84 |
| b (in Å) | 149.84 | 150.86 | 148.84 |
| c (in Å) | 105.49 | 105.61 | 105.34 |
| $\alpha$ (in degrees) | 90 | 90 | 90.00 |
| $\beta$ (in degrees) | 90 | 90 | 90.00 |
| $\gamma$ (in degrees) | 120 | 120 | 120.00 |
| **Resolution (Last shell)** | 40.93-1.85 (1.95-1.85) | 35.23-1.80 (1.92-1.82) | 48.76 -2.35 (2.41-2.35) |
| **Completeness (%)** | 99.1 (100) | 99.3 (100) | 96.24 |
| **<I/$\sigma$ (I)>** | 9.9 (1.79) | 14.0 (3.42) | 8.5 (2.00) |
| **R<sub>merge</sub> (%)** | 13.3 (100) | 14.0 (91.7) | - |
| **Multiplicity** | 11.1 (10.3) | 8.0 (7.4) | - |
| **R-factor (%)** | 18.1 | 16.3 | 16.2 |
| **R-free (%)** | 21.6 | 18.9 | 23.9 |
| **RMS deviations from ideal values** |  |  |  |
| Bond length (Å) | 0.37 | 0.77 | 0.29 |
| Bond angle (°) | 0.56 | 0.85 | 0.40 |
| **Residues (%) in Ramachandran plot** |  |  |  |
| Most favoured regions [A,B,L] | 92.6% | 92% | 94.0% |
| Additional allowed regions [a,b,l,p] | 7.1% | 7.6% | 5.0% |
| Generously allowed regions[~a,~b,~l,~p] | 0.3% | 0.3% | - |
| Disallowed regions [XX] | 0 | 0 | 1% |

AmQNS has a native structure that is similar to other types III PKSs in terms of structural folds (**Figure 4A**), with a unique structural topology that includes a specific upper domain ‘αβαβα’ (ketosynthase domain)^27^, that is conserved in all structural homologs and a lower domain that contains the majority of the substrate-binding residues (A133, E192, T194, T197, S338). In AmQNS monomer, these domains are made up of three beta sheets (13 strands (22.4%), 16 helices (37.6%), 3-10 helices (2.7%), and other secondary structures (37.3%-including four beta hairpins, four beta bulges, 30 beta turns, two gamma turns, and other structures) (**Figure 3, supplementary figure S3**). The protein is functionally active in dimeric form, and in each monomer, the β-sheets are organized into two antiparallel β-sheets and one mixed sheet, where the strands are arranged in the AmQNS structure’s core, whereas the α-helices are distributed on the surface. The prospective substrate-binding pocket entrance of each AmQNS monomeric unit is bordered by the side chains of the α-helices and β-strands. Despite being structurally comparable even at the active site entrance, AmQNS has a considerably bigger binding pocket (volume wise) than its nearest functional homologs from *Citrus X Microcarpa* (PDB IDs: 3WD7 & 3WD8) (**Supplementary figure S4**). This could be the consequence of the single phenylalanine to valine substitution (F265V), where the smaller valine (V) frees up more space in the active site pocket, and the longer tunnel enables the entry of bulky substrates (e.g., *N*-methylanthraniloyl-CoA).

**Figure 4:**
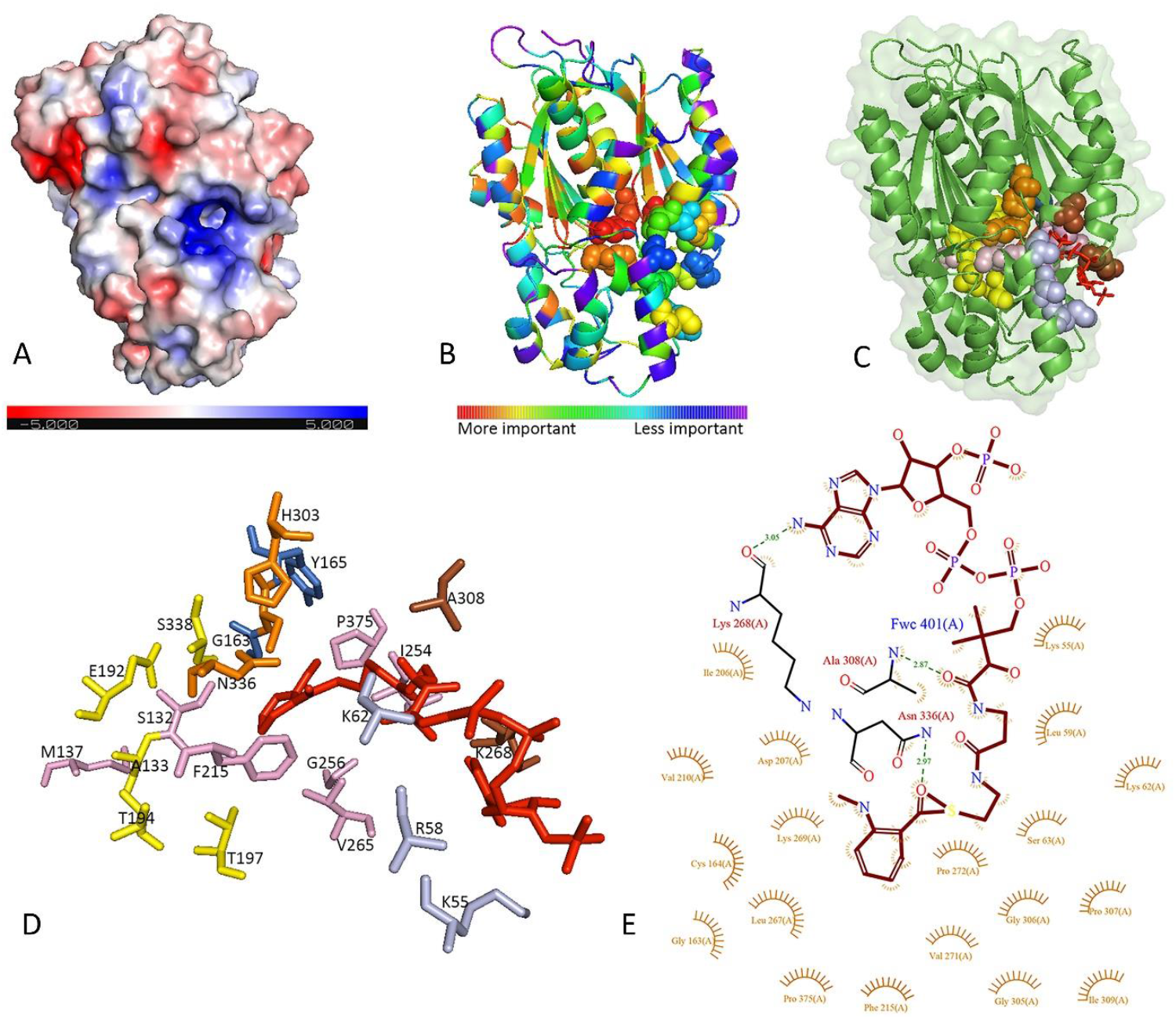
The structure of AmQNS in native and substrate bound form. **A)** AmQNS native structure (surface representation, PDB ID: 6L5U, Resolution 1.85Å) displaying electrostatic charge distribution (positively charged residues in blue and negatively charged residues in red **B)** Evolutionary trace on native AmQNS showing functionally significant residue positions in the structure. **C)** AmQNS-substrate bound form (cartoon and surface representation, PDB ID: 7CCT, Resolution 2.35Å). Red stick represents the substrate *N*-methylanthraniloyl CoA (MANT-CoA). All significant residues are highlighted (catalytic triad (C164, H303, N335)- orange; substrate-binding residues (A133, E192, T194, T197, S338)- yellow; residues in cyclization pocket (S132, M137, F215, I254, G256, V265, P375) -pink; Residues in CoA-binding tunnel (K55, R58, K62)- light blue; residues adjacent to the catalytic C164 (G163, Y165)-marine blue; other significant residues that form polar contacts with substrate (K268, A308)-brown. **D)** Binding environment of MANT-CoA in 7CCT (highlighted the substrate binding area of Figure 4C). MANT-CoA binding pattern (red sticks) and surrounding residues are highlighted. **E)** LIGPLOT of interactions involving ligand MANT-CoA and surrounding residues.

The orientation and locations of the catalytic residues in AmQNS are comparable to those in the closest homologs (**Supplementary Figure S5**). CmACS did not display many differences when analysing the substrate-binding residues of AmQNS. Nonetheless, CmQNS showed modest variations, as demonstrated by changes in the cavity volume metrics (reduced parameters) (**Supplementary Figure S4 & S6**), and even though these sequence alterations and its orientations are minimal, it will result in potential differences in pocket volume, leads to a change in product formation profile. AmQNS prefers bulkier substrates, as previously stated, and the electrostatic potential surface calculation revealed that the substrate-binding pocket regions of AmQNS have a predominantly positive charge (**Figure 4A**), which facilitates binding with the phosphate groups of the preferred starter substrate CoAs. These positions of positive charges are consistently found across type III PKSs as binding of the CoA portion of substrates is conserved. Salt bridges (∼6), hydrogen bonds (∼32), and nonbonded interactions (∼318) connect the hydrophobic and hydrophilic residues that make up the AmQNS dimeric interface area (2463-2481 Å), and salt bridges at the protein interface help to stabilize the protein. The residues involved in forming prospective salt bridges are D96, D136, D251, H257, R259, and K281 from chain A and R259, H257, R146, D136, D96, and E153 from chain B. The catalytic triad (C164-H303-N336) is located in the upper domain, and are deeply embedded within the entrance cavity like their homologs^15^. The orientation and position of these residues are strikingly similar to those of the homologs to some extent. Furthermore, cysteine in the catalytic triad (C164) is highly nucleophilic (as determined by the pKa, due to the reactivity of its thiol (S-H) group), and that this residue is primarily responsible for thioester exchange reactions^28^. The reduction of the sulfur donor molecule in enzyme catalysis is significant since it binds to the substrate, and it is worth noting that ‘thiol groups of cysteines’ are typically found at active sites. This is consistent with prior studies^29^, where the catalytic cysteine in AmQNS is oxidized to sulfinic acid, showing that it has a higher nucleophilicity and is more vulnerable to oxidation^30,31^. Interestingly, molecular evolution also plays a crucial role in maintaining the active-site environment of type III PKS proteins. According to Liou et al.,^31^ CHSs from basal land plants (bryophytes, lycophytes, etc.) have fewer reactive catalytic cysteines than CHSs from higher plants. It is unclear whether these findings regarding the modulation of catalytic cysteine reactivity represents a general pattern in non-chalcone forming PKS family members too. However, AmQNS has a highly nucleophilic cysteine (Cys164) in the catalytic region, indicating that it might have evolved to have a high catalytic potential.

The amino acid residues K268, A308, and N336 form polar contacts (the distance of 3.05Å, 2.87Å, and 2.97Å, respectively) with the CoA molecule in substrate-bound AmQNS (**Figure 4B-D**). The interaction with *N*-methyl anthraniloyl CoA (MANT-CoA) was also confirmed by the presence of 76 nonbonded interactions. One of these residues, N336, is a catalytic site,^32^ and the interactions between all of these residues imply that the main substrate binding sites are in the phosphate region. Likewise, K55, L267, G305, and A308 establish hydrogen bonds (at distances of 2.51Å, 2.93Å, 3.11Å, and 3.06Å, respectively) in the CoASH (byproduct)-bound form (PDB ID: 6L7J, resolution 1.8, **Supplementary Figure S7**). K55 is located in the CoA-binding tunnel at the entrance, and G305 has previously been reported to play a role in shaping the appropriate geometry of the active site pocket^33^ (**Supplementary Figure S7A, B**). Additionally, thermal disorder parameters might indicate conformational flexibility^33^, and we observed that ligand binding causes well-defined conformational changes in proteins, particularly in the β-turn region of AmQNS (residues K268-K269-D270). D270 is absolutely conserved in all aligned proteins, and K269 is mostly conserved (**Figure 2A**), however only AmQNS and CmACS maintain the K268. CmQNS has a K268 to S268 alteration, while other homologs have ‘K268 to L268’. These regions are more flexible, and comparison studies indicated conformational flexibility at the substrate-binding pocket entrance in AmQNS, which suggested hinge-like movement of the surface loop. This flexibility in the AmQNS enzyme structure provides a larger passageway for a substrate to enter the internal active binding site, which is more evident by the followed simulation experiments.

### Structural basis for AmQNS synthetic selectivity

To gain insights into the reaction mechanism, followed by the structural elucidation, molecular simulation studies were used to investigate the mechanistic basis of AmQNS synthetic selectivity. Here we examined if specific ligand-protein interactions can be mapped to characterize the enzyme’s relative propensity to select an optimal number of intermediate ketide insertions. We calculated transition states for MANT-CoA binding to AmQNS and defined the three reaction steps (**Figure 5**) required for AmQNS-driven quinolone production. The first step entails a classic Sn2 thiol addition^34,35^, through which the MANT-CoA substrate binds to the catalytic C164. The second reaction depicts a ketide unit’s concerted process from malonyl-CoA inserts between the cysteine sulfur and the carbonyl carbon of the substrate enzyme complex. The third reaction is then a reverse substitution through which the substrate amine induces product ring closure, which restores the enzymatic cysteine (**Figure 5A**). The chemical structure of MANT-CoA, its derivatives and products (quinolone/ acridones) are given in **Supplementary Figure S8**. The activation energy and enthalpy for each step of the reaction process are provided in **Table 2**.

**Figure 5:**
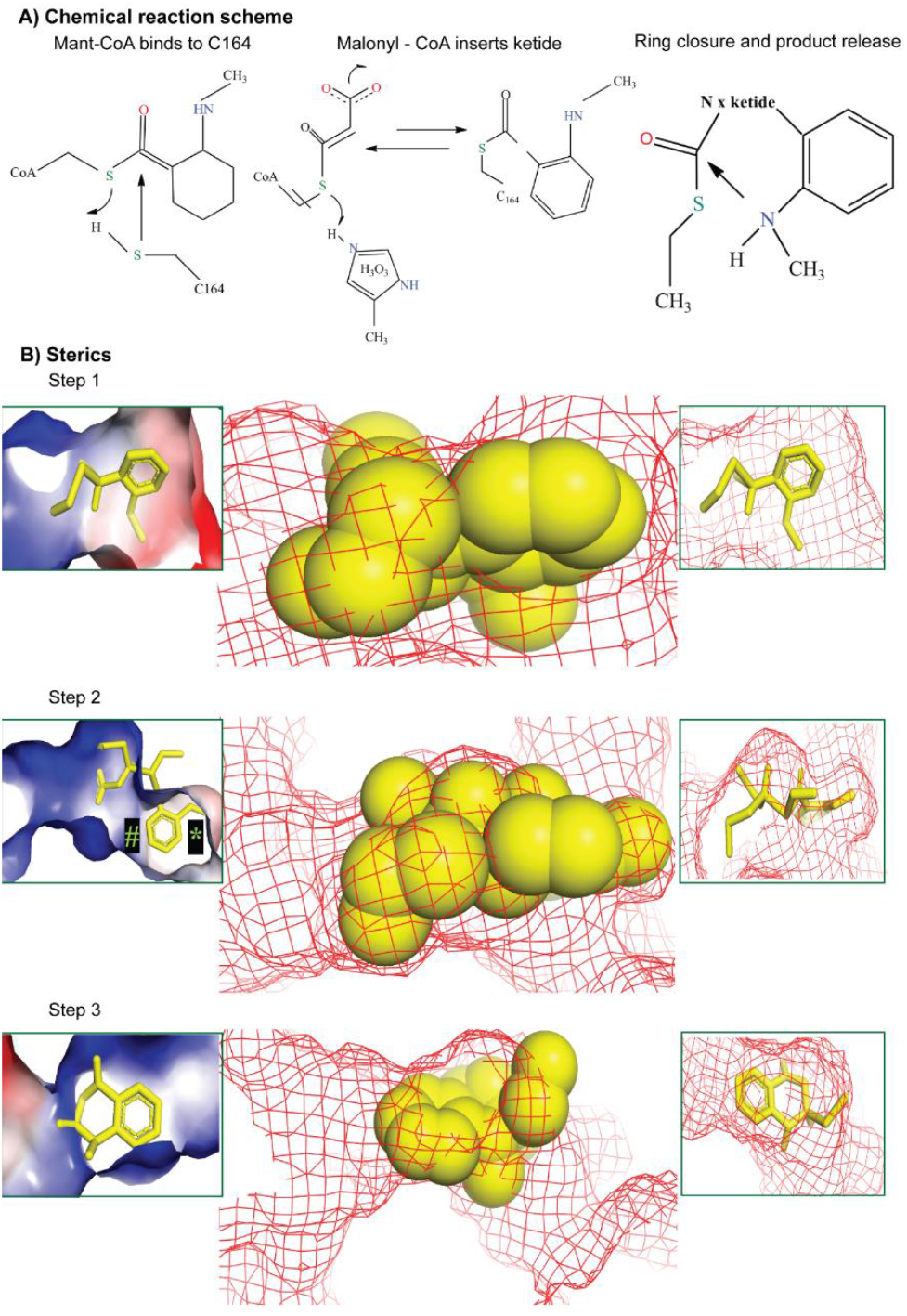
Three steps for AmQNS catalysis. **A)** Chemical reaction scheme showing the substrate binding to the enzyme and subsequent product formation. **B)** Molecular models show different transition state for complexation between the substrate and enzyme. The effect of specific structural features (steric and electrostatics) on kinetic properties for the subsequent reaction (ketide insertion) is represented in the molecular model (‘#’ and ‘*’ markings convey that the receptor poses negligible clashes with the substrate during the first ketide insertion. Still, it should be noted that there is no significant excess of space available at positions ‘#’ and ‘*’ in the first insertion. Consequently, during the process of second and third ketide insertions, during which the reaction intermediate is growing, clashes would be expected at both positions ‘#’ and ‘*.’). Transition steps can be better viewed in **supplementary movies (S1-S3)**.

**Table 2:** MANT quinolone and acridone kinetics.

**Structural dependency for MANT quinolone kinetics and thermodynamics**
|  | 1. Mant-CoA<br>binds C164 | 2. Malonyl-CoA<br>inserts ketide | 3. Quinolone closure<br>& product release |
| --- | --- | --- | --- |
| MANT | Enthalpy: 36.0 kcal/mol<br>Activation: 57.3 kcal/mol | Enthalpy: -3.5 kcal/mol<br>Activation: 39.6 kcal/mol | Enthalpy: -5.3 kcal/mol<br>Activation: 15.1 kcal/mol |
| MANT-pyridine | Enthalpy: 40.4 kcal/mol<br>Activation: 60.2 kcal/mol | Enthalpy: -9.2 kcal/mol<br>Activation: 31.4 kcal/mol | Enthalpy -4.1 kcal/mol<br>Activation: 23.0 kcal/mol |
| MANT-chloro | Enthalpy: 32.7 kcal/mol<br>Activation: 53.5 kcal/mol | Enthalpy: 4.4 kcal/mol<br>Activation: 44.1 kcal/mol | Enthalpy: -4.6 kcal/mol<br>Activation: 19.9 kcal/mol |

**Structural dependency for MANT acridone kinetics and thermodynamics**
|  | 1. 2nd Malonyl-CoA<br>inserts ketide | 2. 3rd Malonyl-CoA<br>inserts ketide | 3. Acridone closure and<br>product release |
| --- | --- | --- | --- |
|  | Enthalpy: 10.4 kcal/mol<br>Activation: 59.9 kcal/mol | Enthalpy: 9.4 kcal/mol<br>Activation: 53 kcal/mol | Enthalpy: -16.0 kcal/mol<br>Activation: 38.3 kcal/mol |

The multiple transition states^36–39^ for initial complexation between the substrate and enzyme were then demonstrated (**Figure 5B**). Transition states tend to be the portion of any reactive progression, where structural features have the most significant influence on kinetic properties for the subsequent reaction. In the particular case of MANT-CoA binding to the AmQNS active site, it is apparent that the substrate is a good fit for the enzyme, as there are minimal clashes that could either kinetically disfavour or completely abrogate the subsequent reaction. Several observations are made regarding areas on the substrate for which clash is so minimal that (theoretically) increased substrate bulk might reduce the activation barrier (i.e., improve reaction kinetics) through favourable van der Waals (vdW) or electrostatic interactions^40–42^. The observation that the receptor is spacious around the aminomethyl substrate group led to the notion of experimentally investigating whether (computationally) the aminomethyl group could be productively modified as a chloro analog (somewhat bulkier in a potentially favourable manner). Notably, the red and blue receptor patches in the MANT aryl ring region are similarly motivated to explore if a slightly more polar version of the substrate (with a pyridinyl ring, rather than benzyl) might produce kinetically favourable electrostatic complementarity.

Notably, step 2 in the reaction sterics (**Figure 5B**) represents prospective receptor structural influences on the first ketide insertion kinetics. In this case, the transition state for this reaction is well accommodated by the receptor, which, in turn, corroborates the prior observation that AmQNS is a viable ‘enzymatic engine’ for promoting quinolone synthesis. Importantly, we propose that second and third ketide insertions may be somewhat less favored than the first insertion. Nonetheless, during the first insertion, there is no considerable excess of space available. As a result, clashes would be expected (in places ‘#’ and ‘*’) during the second and third ketide insertions, when the reaction intermediate is growing. These clashes may be somewhat defused with a ligand conformational shift that orients the ring slightly out of the plane of this graphic as the aryl ring begins to progress toward the narrow product exit channel, whose position is relatively well marked in the figure (‘*’). In step 3, we see transitional interactions between the forming quinolone product and the receptor. It is interesting to note that although the receptor is not hugely antagonistic to product formation, it also does not seem ideally suited, as apparent in the steric clash between the enzymatic surface and the aminomethyl. This clash might be alleviated through a change of conformational twist (to reorient the aryl ring) that is essentially the same factor identified earlier in reaction step 2 as a requisite step for second or third ketide insertions. This has an exciting implication and, this means that although the analysis of step 2 has pointed firmly toward smaller quinolone product formation (compared to a larger acridone product), a kinetic hitch in the final step of quinolone formation may nullify this difference. We suggest that structural modifications to the AmQNS enzyme (e.g., potentially mutating Leu 263 into a smaller valine or alanine or removing the methylamine clash by mutating Ser 132 into a glycine) might favour both the quinolone and acridone product formation, potentially speeding the production of either while not necessarily affecting the relative ratios of quinolone and acridone product. In **Table 2**, we show the computed impacts of the two minor (chloro and pyridinyl) modifications to the MANT-CoA substrate. Our data show that the substrate modifications appear to have only minor influence, and it is difficult to predict if either shift will produce a demonstrable improvement in reactive profile relative to unmodified MANT-CoA. Alternatively, we also suggest that AmQNS may support the catalytic a variety analogs to the standard biologically processed substrates, meaning that their synthetic chemistry can be extended from the production of novel natural product scaffolds to a related display chemical analogs.

Next, we report acridone-specific reaction steps (**Figure 6A**) and the second and third ketide insertions are predicted to be somewhat less favourable kinetically and thermodynamically compared to the first ketide insertion shown in **figure 5**. In contrast, the final acridone ring closure is expected to have a higher activation barrier than the quinolone product formation but a more favourable reaction enthalpy. Finally, we investigated whether AmQNS is better suited for quinolone or acridone production, and we propose that the key difference between the two reactions may be a matter of stoichiometric control, with an excess of malonyl-CoA favoring acridone and tight stoichiometry favoring quinolone. Furthermore, reducing steric bulk by altering Leu 263 or Ser 132 could enhance throughput of both products, indicating that specific amino acid changes could be used to impact enzymatic product selectivity. For instance, the previously studied^12^ AmQNS mutants MSD1 (double mutant, S132T/A133S) and MSD2 (triple mutant, S132T/A133S/V265F) had drastically narrowed active site cavities when compared to the wild-type AmQNS. MSD1 demonstrated chalcone-forming activity with p-coumaroyl-CoA like the typical chalcone synthase, whereas MSD2 did not^12^. Since none of the mutants prefer MANT-CoA as starter substrate, the two amino acid alterations S132T and A133S influenced the enzyme’s substrate selectivity.

**Figure 6:**
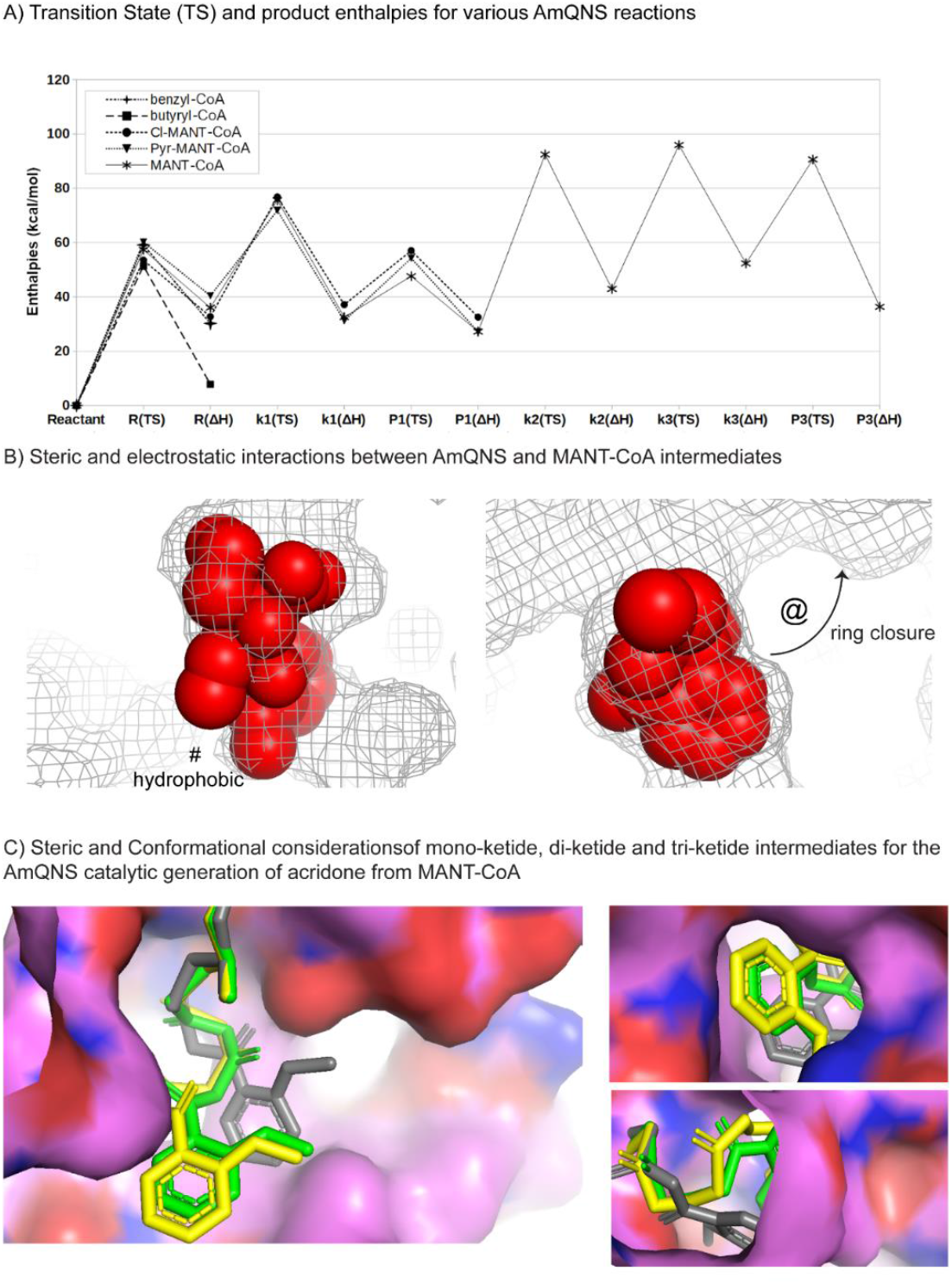
AmQNS mediated synthetic diversity based on the structure conformations. **A)** Relative transition state (TS) and product enthalpies for various AmQNS reactions, relating to five distinct core substrates named in the legend (upper left; Cl-MANT refers to a chloromethyl analog to MANT; Pyr-MANT refers to a pyridinyl analog to MANT, with the heteroatomic N located para to the methylamine). Specific reaction state enthalpies are quantified for initial binding (R), specific polyketide insertions (k1, k2, k3) and product cleavages after 1st (P1) and third (P3) ketide insertions. **B)** Steric and electrostatic interactions between the AmQNS receptor (mesh) and MANT-CoA-based intermediates (spheres) for the ketide insertion (shown in left) and monoketide product forming (shown in the right) transition states. Steric effects relating to the bulky chlorine atom are represented as ‘@.’ **‘**#’ indicating the hydrophobic pocket. **C)** Steric and conformational considerations of mono-ketide (yellow), di-ketide (pink) and tri-ketide (white) intermediates for the AmQNS catalytic generation of acridone from MANT-CoA substrate, as viewed from a cross-section of the whole receptor, the channel through which malonyl-CoA co=substrate enters, and the product release channel.

It is inherently challenging to fully characterize how type III polyketide quinolone synthases achieve such impressive synthetic diversity from relatively minor structural variations among different enzyme families. Here, we primarily focus on enzymatic steric and electrostatic selectivity for binding other core substrate units (e.g., relative favorability for specialized binding units such as coumaroyl-CoA, benzyl-CoA, acetyl-CoA, MANT-CoA, versus a universal capacity to malonyl-CoA as a substrate or co-substrate), and the amount of space available to accommodate larger numbers of incrementally inserted ketide units. We determined quantum chemical transition states to compare the relative kinetic barriers^43^ and thermodynamic enthalpies^44^ for the initial complexation of MANT-CoA, benzyl-CoA, butyryl-CoA and coumaroyl-CoA. Similar characterization was done, in the case of MANT-based reactions, for the first ketide insertion, the single-ketide quinolone product formation, the second and third ketide insertions, and the triple-ketide acridone product.

The initial complexation barrier showed little variation among primary substrates, implying that steric dependencies play a minor impact at this stage (**Figure 6A**). However, steric and electrostatics do appear to discriminate somewhat in the stability of the resulting bound intermediates. Specifically, the pyridinyl analog to MANT-CoA has less stabilization than the others because it places the slightly polar aryl nitrogen directly within a hydrophobic pocket delimited by Ile 254 and Pro 375 (‘#’ in **Figure 6B**). Simultaneously, the sole flexible substrate (butyryl-CoA) can conformationally adapt to this pocket in a stabilizing manner. Proceeding from the mono-ketide intermediate (k1) to the monoketide product (P1) reflects minimal difference among the three analogs of MANT-CoA, with the exception that the chloromethyl compound has a higher barrier to ring closure, due to steric effects relating to the bulky chlorine atom.

Next, in the **Figure 6A**, it showed the quantitative reaction profile of secondary and tertiary ketide insertions by which the monoketide intermediate may progress toward the acridone product (P3). It is worth noting that these latter ketide insertions are predicted to have higher barriers (> 50 kcal/mol) than the first insertion (∼40 kcal/mol). From a computational perspective, this trend is rationalized by higher conformational strain evident in the di-ketide and tri-ketide units relative to the mono-ketide (**Figure 5B**), as opposed to issues relating to the approach of malonyl-CoA co-substrate (for which there is ample space, as shown in Figure 6C) or the situation of the MANT group (**Figure 5B**). Based on this, we propose that AmQNS structurally favours the smaller quinolone product’s production and might thus only produce acridone under conditions of significant malonyl-CoA concentration.

In the future, we will compare analyses for non-MANT substrates. Furthermore, the specific structures available for each of the five (for quinolone) transition states or five (for acridone) reaction steps may be rigorously evaluated to determine which ligand-receptor amino acid contacts are favourable or unfavourable. Also, determine what sorts of mutations could be proposed to significantly alter favourability in a manner that could influence existing AmQNS enzymatic activity and potentially engineer product specificity

## Materials and Methods

### Homolog identification and phylogenetic analysis

To identify the potential homologs for AmQNS, complete predicted proteome sequences were retrieved from JGI (http://genome.jgi.doe.gov) and NCBI GeneBank (https://ncbi.nlm.nih.gov/). The Marine Microbial Eukaryote Transcriptome Sequencing Project database (MMETSP)^45^ were also used to predict the homologs. The *Arabidopsis thaliana* chalcone synthase (AT5G13930) reference amino acid sequence was used as a query to search of all potential homologs using the Hidden Markov model (HMM)-based tool jackhammer^46^. Evolutionary genealogy of genes: Non-supervised Orthologous Groups (eggNOG) mapper was used for hierarchical resolution of orthology assignments^47^. Finally, the SMART and Pfam databases were employed to identify conserved domains present in type III PKS from different organisms^48,49^, both SMART and Pfam databases were merged, and redundant domains were filtered-out and the Hidden Markov model (HMM)-based tool hmmscan (https://github.com/EddyRivasLab/hmmer). Only sequences with the catalytic or conserved domain of the references were retained. The identified homologs and the respective details are given in the **supplementary table S1**. All identified homologs were aligned using MAFFT^50^ and ambiguously aligned regions were excluded for further analysis using trimAl software^51^. Alignments were tested using ProtTest v3^52^ to choose an appropriate model for nucleotide substitution. Two separated Maximum likelihood (ML) phylogenetic trees were computed using RAxML-NG^53^ and IQ-TREE2^54^. ML analyses were performed using 1000 bootstrap replicates. The supporting values from both software were noted on the ML-unrooted tree.

### Large-scale AmQNS Expression and Purification

The QNS gene from ‘*A. marmelos*’ was cloned into pET32b as explained by Resmi et al., 2013^12^. Transformed *Escherichia coli* BL21(DE3) cells were incubated at 37°C in Luria-Bertani (LB) medium containing ampicillin (100 g/ml) until they reached the exponential phase of growth (OD600 0.6). Isopropyl 1-thio—D-galactopyranoside (IPTG, 400 mM) was employed to induce AmQNS expression, and the cells were further incubated at 28°C for 5-6 hours. At 4°C, all phases of protein purification were carried out. The pellet was resuspended in KPO4 buffer (50 MM, pH 8) containing NaCl (0.1 M), imidazole (40 mM), and lysozyme (750 μg/ml) after centrifugation (5000 g, 30 minutes, 4°C). The lysate was sonicated (amplitude: 35%, 3 seconds on, 5 seconds off, 30 minutes) on ice after being incubated for half an hour on ice. The lysate was then centrifuged (10,000 g, 60 minutes) and the supernatant was then loaded to a Ni-NTA (nickel-nitrilotriacetic acid) affinity column equilibrated with KPO4 buffer (50 mM, pH 7.9) containing NaCl (0.5M) and imidazole (40 mM). In the resuspended condition, the system was allowed to bind at 4°C (1-2 hours). The recombinant protein was eluted in 15 mM KPO4 (pH 7.5) buffer containing 500 mM NaCl, 500 MM imidazole, and 10% glycerol after a lengthy wash of the column with the same equilibration buffer (10 column volumes). Purified recombinant AmQNS (61 kDa) fractions (containing an N-terminal Trx-S-His fusion tag) were concentrated (Amicon-Ultra centrifugal filters, 10 kDa cut-off) and overnight enterokinase cleavage was performed to remove the fusion tag1. Size exclusion chromatography on a Superdex 200HR (10/100 GL) column (GE Healthcare) in HEPES-NaOH buffer (20 mM, pH 7.5) containing NaCl (100 mM) and dithiothreitol was used to further purify the AmQNS protein solution to homogeneity (a monomeric molecular weight of 43 kDa) (DTT, 2 mM). The purified AmQNS fractions were further concentrated to 20 mg/ml in the same HEPES. SDS-PAGE was used for qualitative analysis, and the quantity was also calculated using the NanoDropTM1000 spectrophotometer (Thermo Scientific, Wilmington, DE) at an optical density (OD) ratio of 260/280 (and default protein absorbance values for 0.1%. i.e., 1 mg/mL). MALDI TOF MS analysis was used to determine the protein’s homogeneity and mass accuracy. The monomeric molecular weight of the resulting protein was 43 kDa, which was consistent with the expected molecular mass of 43.8 Da (the calculated molecular mass of AmQNS is 42.8 kDa, with the inclusion of approximately 10 amino acid residues from the vector causing the change in molecular mass). The molecular weight of the purified protein obtained by MALDI is 43.9 kDa (data not shown) and corresponds roughly to the expected molecular weight of the full-length polypeptide chain.

### AmQNS Crystallization by microbatch method

Diffraction quality crystals were obtained in both native and substrate bound forms and optimized the conditions. AmQNS could produce a good diffraction quality crystal which diffracted up to 1.85 Å (H32 space group), when using 1:1 drop ratio by using 2 *µ*L of protein and 2 *µ*L of precipitant (0.1 M HEPES 7.5, 1.4M Sodium citrate tribasic; Index 20 of Hamptons research screen) with additives (0.1M Magnesium chloride hexahydrate or 0.1M Cadmium chloride hydrate). Co-crystallization trials were performed in presence of its favourable CoA substrates ‘N-methyl anthraniloyl CoA (MANT-CoA)’. Solutions containing the substrate was directly added to the concentrated protein solution to a final concentration of ∼2mM and incubated in ice for an hour prior to crystallization experiments. Co-crystallization studies were conducted by microbatch method. Diffraction quality of these crystals was also optimized wherever necessary by adding additives, varying drop sizes and protein/precipitant concentrations. The crystals appeared within a span of 2-3 weeks with approximate dimensions of 0.1 mm X 0.1 mm X 0.1 mm. Soaking experiments were also conducted in addition to co-crystallization experiment. Soaking native crystals with substrates is often the method to obtain crystals of protein–ligand complexes. Here, AmQNS native crystal was soaked (30 minutes) in ligand solution, which was prepared in the same crystallization condition. I.e., 2mM *N*-methylanthraniloyl-CoA was used for AmQNS-substrate soaking experiments (Index 20+0.1M Cadmium chloride hydrate). We could also get the byproduct (CoASH) bound AmQNS crystals. After proper incubation, crystal was picked with a nylon loop, flash frozen in liquid nitrogen and data were collected.

### X-ray Data Collection, processing, and Refinement

After obtaining diffraction-quality crystals, the crystals were cryoprotected (20% glycerol in condition buffer/crystallization solution) by plunging them into liquid nitrogen using a fine-gauge wire micro loop. Data from the native and CoASH bound crystals belonging to the space group H32 were collected at the Molecular Biophysics Unit (IISc, Bangalore) using a MAR 345 image-plate detector mounted on a Bruker MICROSTAR ULTRA II Cu Kα rotating anode X-ray generator (wavelength of 1.54179 Å). For collecting the high-resolution data, the spacing between the protein crystal and the detector was adjusted to 200 mm. All data were collected at 100 K. Data collection statistics are given in **Table S1**. iMosflm was used to process the diffraction images^55^, and data were merged using SCALA^56^ in the CCP4^57^. The intensity data were converted into structure-factor amplitudes using TRUNCATE in the CCP4^58,59^.

Further, structures were solved by the molecular replacement method at 1.85Ǻ, 2.35Ǻ for the H32 space groups. The structure of AmQNS co-crystallized with *N*-methylanthraniloyl-CoA was solved at a resolution of 2.35 Ǻ. PHASER^**60**^ in the CCP4 suite^**57**^ was used for molecular replacement by employing the structure of the acridone synthase from *Citrus microcarpa* (PDB ID: WD7; identity-93%) as the search model^**15**^. The solutions obtained from molecular replacement were subsequently refined using REFMAC5^**61**^, along with multiple rounds of manual model building using COOT v0.7.1^**62**^,^**63**^. Addition of the ligands and water atoms was performed by PRODRG^**64**^. The possibility of alternate ligand conformations was also evaluated before finalizing the ligand fitting. The final refinement of the native structures was performed in PHENIX^**65**^. Images of the protein structures were generated using PyMOL Licenced academic version^**66**^. The refined models were validated by PROCHECK^**67**^ and the MOLPROBITY^**68**^. All structural models were manually built, refined, and rebuilt with REFMAC5/PHENIX and COOT.

### Structural Analysis

The refined protein structures were evaluated using MolProbity with the Phoenix server and wwPDB server^69^. Structural alignments were performed in ALIGN (Pymol^66^) and mTM-align^70^. The sequence-structure conservation patterns were analysed using ESPript and ENDscript 2.0^24^. The neighbour-joining method^71^ was used to construct a structural phylogram. 2F0-Fc maps were calculated in CCP4 v7.0 using the ‘fft’ module, and the maps were visualized in PyMOL using the command line option (contoured at 1.0 sigma around the selection site within 1.6 Å of the selected atoms). The electrostatic properties of AmQNS were calculated using APBS using the PyMOL plugin. PDB2PQR Version 2.0.0^72^ was used to convert the PDB files into PQR files. To obtain the detailed characteristic features of the surface pockets and interior voids of AmQNS, CASTp (Computed Atlas of the Surface Topography of Proteins) was used^73^. The default probe radius was used (1.4 Å). The protein secondary structure and protein-ligand interactions (determined using LIGPLOT) were analysed using PDBsum (www.ebi.ac.uk/pdbsum) ^74^. The relative position of functional and structural importance among the protein homolog sequence sites was estimated using Evolutionary Trace (ET; http://evolution.lichtargelab.org/). All figures were prepared using PyMOL v2.4.1^66^.

### Surface Plasmon Resonance (SPR) based assays

#### AmQNS immobilization on sensor chip

*ProteOn* ‘GLM’ chip was used for the SPR interaction studies where AmQNS could give good response (L3 - 9962 RU, L4 - ∼7000) on immobilization (Supplementary **Figure S2**). Here, amine coupling works where the amine groups present in the AmQNS covalently bind to chemically activated carboxyl groups of the dextran molecules. Channel L3 and L4 were used for immobilization of protein while L2 was used as reference. The extent of non-specific interactions was eliminated or reduced by optimizing the buffer conditions. Furthermore, ligand stability on the GLM biosensor over time was checked over a period of 30 days and was found that it is active, by resulting in quantifiable interactions. This demonstrates that the AmQNS immobilization onto the GLM sensor surface does not limit the functionality, confirming the use of this SPR label-free technology to study its interaction pattern with different acyl-CoA substrates. *N*-methylanthraniloyl-CoA was purchased from TransMIT (Plant MetaChem, www.plantmetachem.com), whereas all other substrates were purchased from Sigma-Aldrich (www.sigmaaldrich.com)

#### Binding kinetics studies of AmQNS with different substrates

All binding studies were performed at 30°C in ProteOn XPR array system^75^. The SPR based system measures the changes in refractive index to investigate the direct interaction between AmQNS and different CoA substrates. The analytes (substrates) were injected over the surface of the chip and any binding between the two resulted in the change in surface mass, which is recorded, and measures as a change in refractive index. In our experiments, AmQNS was captured on the surface of the GLM sensor chip and used to screen the preferred substrates (acyl-CoA’s) in the presence and absence of malonyl-CoA. We could not find any interaction in the absence of malonyl-CoA, and it was quite interesting to note that the binding modes of these CoA substrates to AmQNS are influenced by the presence of the malonyl-CoA, which is the extender during the biochemical reaction of polyketide formation. Sensogram prepared by processing the data (after subtraction of L2 responses (reference channel). Baseline drift due to the bulk refractive index change, non-specific binding, matrix effects and injection noise were also corrected using the reference spots. Further, the responses obtained from the AmQNS-small molecular interactions at different concentrations were fitted using the Langmuir 1:1 biomolecular interaction model using the ProteOn Manager software version 3.1.0.6 (Bio-Rad, USA). Equilibrium dissociation constants (*K*D) were calculated from the ratio of the association and dissociation rates.

### Molecular simulation studies

The AmQNS polyketide synthase active site structural model was constructed from an AmQNS crystallographic model (PDB ID: 6L5U) in PyMol^76^ as the set of all amino acids with at least one atom residing within 12.0 Å of the catalytic Cysteine (C164). Peripheral peptide chain termini were neutralized by simple protonation to neutral amine and aldehyde structures. Specific ligands (MANT-CoA and malonyl-CoA) were constructed *in situ* using PyMol by referring to the -EthSH group of the co-crystallized CoASH ligand. For computational efficiency, the bulk of the conserved CoA moiety (i.e., all except for those mentioned above - EthSH moiety) was removed from each ligand. Transition state calculations were performed using MOPAC 2016 ^77^, via the PM7 parametrization ^78,79^. Due to the exceptional complexity in the potential energy surface (PES)^80,81^, it was necessary to manually perform transition states by employing constraints to implement a stepwise approach between reacting atoms.

We also calculated transition states for MANT-CoA binding to AmQNS. An initial step size of 0.4 Å was used for the initial (distant) ligand approach until the approaching atoms were within 1.0 Å of the expected covalent distance. A step size of 0.1 Å was employed to capture subtle structural and energetic effects. All receptor backbone atoms were held rigid to prevent spurious peripheral conformational shifts from quantitatively overwhelming covalent energetics, as were all side chains except those participating directly in enzyme reaction function. In addition, we also determined quantum chemical transition states to compare the relative kinetic barriers and thermodynamic enthalpies for the initial complexation of MANT-CoA, benzyl-CoA, butyryl-CoA and coumaroyl-CoA.

### Accession codes

Coordinates and structure factors for the above mentioned AmQNS structures have been deposited in the Protein Data Bank (accession codes: 6L5U, 6L7J and, 7CCT).

## Acknowledgments

We are grateful to Prof. Dr. M. Vijayan and Dr. B. Gopal, Molecular Biophysics Unit, Indian Institute of Science, Bangalore, to use the X-ray lab facilities. We thank Arun Surendran for SPR technical support and Dr. Abdul Jaleel for allowing to use the proteomics facility. The authors also acknowledge Devika Vikraman for assisting in figure preparation. MV acknowledges the Council of Scientific and Industrial Research (CSIR) for Research Associateship (09/716(0178)/2018-EMR-1 dated 26.04.2018) and the Department of Biotechnology (DBT) for financial support.

## Author Contribution

EVS supervised the study. MV contributed to conceptualization, investigation, methodology, visualization and writing—original draft, review, and editing. AKV contributed to writing, helped in data collection, structure solutions and calculations. DB helped in structure related calculations and critically evaluated the manuscript. KRM assisted in simulation studies and writing. AS helped for phylogeny and contributed to writing. MRP provided scientific advice and provided constant help throughout the studies. All authors have read and agreed to the published version of the manuscript.

## References

1. Shang, S. & Tan, D. S. Advancing chemistry and biology through diversity-oriented synthesis of natural product-like libraries. Current Opinion in Chemical Biology 9, 248–258 (2005).

2. Girija, A. et al. Harnessing the natural pool of polyketide and non-ribosomal peptide family: A route map towards novel drug development. Current Molecular Pharmacology 14, (2021).

3. Shimizu, Y., Ogata, H. & Goto, S. Type III Polyketide Synthases: Functional Classification and Phylogenomics. ChemBioChem 18, 50–65 (2017).

4. Austin, M. B. & Noel, J. P. The chalcone synthase superfamily of type III polyketide synthases. Natural Product Reports 20, 79–110 (2003).

5. Shang, S., Iwadare, H., Macks, D. E., Ambrosini, L. M. & Tan, D. S. A Unified Synthetic Approach to Polyketides Having Both Skeletal and Stereochemical Diversity. Organic Letters 9, 1895–1898 (2007).

6. Satou, R. et al. Structural basis for cyclization specificity of two Azotobacter type III polyketide synthases: a single amino acid substitution reverses their cyclization specificity. The Journal of biological chemistry 288, 34146–34157 (2013).

7. Austin, M. B. et al. Crystal Structure of a Bacterial Type III Polyketide Synthase and Enzymatic Control of Reactive Polyketide Intermediates. Journal of Biological Chemistry 279, 45162–45174 (2004).

8. Pandith, S. A., Ramazan, S., Khan, M. I., Reshi, Z. A. & Shah, M. A. Chalcone synthases (CHSs): the symbolic type III polyketide synthases. Planta 251, 15 (2019).

9. Hashimoto, M., Nonaka, T. & Fujii, I. Fungal type III polyketide synthases. Natural Product Reports 31, 1306–1317 (2014).

10. Schröder, G. & Schröder, G. Stilbene and Chalcone Synthases: Related Enzymes with Key Functions in Plant-Specific Pathways. Zeitschrift für Naturforschung C 45, 1–8 (1990).

11. Koskela, S., Elomaa, P., Helariutta, Y., Söderholm, P. & Vuorela, P. TWO BIOACTIVE COMPOUNDS AND A NOVEL CHALCONE SYNTHASELIKE ENZYME IDENTIFIED IN GERBERA HYBRIDA. in Acta Horticulturae 271–274 (International Society for Horticultural Science (ISHS), Leuven, Belgium, 2001). doi:10.17660/ActaHortic.2001.560.52.

12. Resmi, M. S., Verma, P., Gokhale, R. S. & Soniya, E. V. Identification and characterization of a type III polyketide synthase involved in quinolone alkaloid biosynthesis from Aegle marmelos Correa. The Journal of biological chemistry 288, 7271–7281 (2013).

13. Heeb, S. et al. Quinolones: from antibiotics to autoinducers. FEMS microbiology reviews 35, 247–274 (2011).

14. Ahmed, A. & Daneshtalab, M. Nonclassical biological activities of quinolone derivatives. Journal of pharmacy & pharmaceutical sciences: a publication of the Canadian Society for Pharmaceutical Sciences, Societe canadienne des sciences pharmaceutiques 15, 52–72 (2012).

15. Mori, T. et al. Cloning and structure-function analyses of quinolone-and acridone-producing novel type III polyketide synthases from Citrus microcarpa. The Journal of biological chemistry 288, 28845–28858 (2013).

16. Navarro-Muñoz, J. C. & Collemare, J. Evolutionary Histories of Type III Polyketide Synthases in Fungi. Frontiers in Microbiology 10, (2020).

17. De Luca, D. & Lauritano, C. In Silico Identification of Type III PKS Chalcone and Stilbene Synthase Homologs in Marine Photosynthetic Organisms. Biology 9, (2020).

18. Zaremba-Niedzwiedzka, K. et al. Asgard archaea illuminate the origin of eukaryotic cellular complexity. Nature 541, 353–358 (2017).

19. Eme, L., Spang, A., Lombard, J., Stairs, C. W. & Ettema, T. J. G. Archaea and the origin of eukaryotes. Nature Reviews Microbiology 15, 711–723 (2017).

20. Mallika, V., Sivakumar, K. C., Aiswarya, G. & Soniya, E. V. In silico approaches illustrate the evolutionary pattern and protein-small molecule interactions of quinolone synthase from Aegle marmelos Correa. Journal of Biomolecular Structure and Dynamics 37, 195–209 (2019).

21. Naake, T., Maeda, H. A., Proost, S., Tohge, T. & Fernie, A. R. Kingdom-wide analysis of the evolution of the plant type III polyketide synthase superfamily. Plant Physiology 185, 857–875 (2021).

22. Stairs, C. W. et al. Chlamydial contribution to anaerobic metabolism during eukaryotic evolution. Science Advances 6, eabb7258.

23. Larkin, M. A. et al. Clustal W and Clustal X version 2.0. Bioinformatics 23, 2947–2948 (2007).

24. Robert, X. & Gouet, P. Deciphering key features in protein structures with the new ENDscript server. Nucleic Acids Research 42, W320–W324 (2014).

25. Bronner, V., Bravman, T., Nimri, S. & Lavie, K. Rapid and efficient determination of kinetic rate constants using the ProteOn XPR36 protein interaction array system. Bio-Rad bulletin 3172 (2006).

26. Goodsell, D. S. et al. RCSB Protein Data Bank: Enabling biomedical research and drug discovery. Protein Science 29, 52–65 (2020).

27. Bräuer, A. et al. Structural snapshots of the minimal PKS system responsible for octaketide biosynthesis. Nature Chemistry 12, 755–763 (2020).

28. Jez, J. M. & Noel, J. P. Mechanism of Chalcone Synthase: pKa OF THE CATALYTIC CYSTEINE AND THE ROLE OF THE CONSERVED HISTIDINE IN A PLANT POLYKETIDE SYNTHASE*. Journal of Biological Chemistry 275, 39640–39646 (2000).

29. Richau, K. H. et al. Subclassification and Biochemical Analysis of Plant Papain-Like Cysteine Proteases Displays Subfamily-Specific Characteristics. Plant Physiology 158, 1583 (2012).

30. Tseng, C. C., McLoughlin, S. M., Kelleher, N. L. & Walsh, C. T. Role of the Active Site Cysteine of DpgA, a Bacterial Type III Polyketide Synthase. Biochemistry 43, 970–980 (2004).

31. Liou, G., Chiang, Y.-C., Wang, Y. & Weng, J.-K. Mechanistic basis for the evolution of chalcone synthase catalytic cysteine reactivity in land plants. The Journal of biological chemistry 293, 18601–18612 (2018).

32. Jez, J. M., Ferrer, J.-L., Bowman, M. E., Dixon, R. A. & Noel, J. P. Dissection of Malonyl-Coenzyme A Decarboxylation from Polyketide Formation in the Reaction Mechanism of a Plant Polyketide Synthase. Biochemistry 39, 890–902 (2000).

33. Wani, T. A. et al. Molecular and functional characterization of two isoforms of chalcone synthase and their expression analysis in relation to flavonoid constituents in Grewia asiatica L. PLOS ONE 12, e0179155.(2017).

34. Hamlin, T. A., Swart, M. & Bickelhaupt, F. M. Nucleophilic Substitution (SN2): Dependence on Nucleophile, Leaving Group, Central Atom, Substituents, and Solvent. ChemPhysChem 19, 1315–1330 (2018).

35. Leichert, L. I. & Jakob, U. Protein thiol modifications visualized in vivo. PLoS biology 2, e333–e333 (2004).

36. Schramm, V. L. Enzymatic Transition States and Drug Design. Chemical reviews 118, 11194–11258 (2018).

37. Roston, D. & Cui, Q. QM/MM Analysis of Transition States and Transition State Analogues in Metalloenzymes. Methods in enzymology 577, 213–250 (2016).

38. Lundberg, M., Kawatsu, T., Vreven, T., Frisch, M. J. & Morokuma, K. Transition States in a Protein Environment −ONIOM QM:MM Modeling of Isopenicillin N Synthesis. Journal of Chemical Theory and Computation 5, 222–234 (2009).

39. Grambow, C. A., Pattanaik, L. & Green, W. H. Reactants, products, and transition states of elementary chemical reactions based on quantum chemistry. Scientific Data 7, 137 (2020).

40. Rifai, E. A., van Dijk, M., Vermeulen, N. P. E., Yanuar, A. & Geerke, D. P. A Comparative Linear Interaction Energy and MM/PBSA Study on SIRT1–Ligand Binding Free Energy Calculation. Journal of Chemical Information and Modeling 59, 4018–4033 (2019).

41. Ren, P. et al. Biomolecular electrostatics and solvation: a computational perspective. Quarterly reviews of biophysics 45, 427–491 (2012).

42. Cruz, J. N. et al. Molecular dynamics simulation and binding free energy studies of novel leads belonging to the benzofuran class inhibitors of Mycobacterium tuberculosis Polyketide Synthase 13. Journal of Biomolecular Structure and Dynamics 37, 1616–1627 (2019).

43. Bernetti, M., Cavalli, A. & Mollica, L. Protein-ligand (un)binding kinetics as a new paradigm for drug discovery at the crossroad between experiments and modelling. MedChemComm 8, 534–550 (2017).

44. Du, X. et al. Insights into Protein-Ligand Interactions: Mechanisms, Models, and Methods. International journal of molecular sciences 17, 144 (2016).

45. Keeling, P. J. et al. The Marine Microbial Eukaryote Transcriptome Sequencing Project (MMETSP): Illuminating the Functional Diversity of Eukaryotic Life in the Oceans through Transcriptome Sequencing. PLoS Biology (2014) doi:10.1371/journal.pbio.1001889.

46. Johnson, L. S., Eddy, S. R. & Portugaly, E. Hidden Markov model speed heuristic and iterative HMM search procedure. BMC Bioinformatics 11, 431 (2010).

47. Huerta-Cepas, J. et al. EggNOG 5.0: A hierarchical, functionally and phylogenetically annotated orthology resource based on 5090 organisms and 2502 viruses. Nucleic Acids Research (2019) doi:10.1093/nar/gky1085.

48. El-Gebali, S. et al. The Pfam protein families database in 2019. Nucleic Acids Research (2019) doi:10.1093/nar/gky995.

49. Letunic, I. & Bork, P. 20 years of the SMART protein domain annotation resource. Nucleic Acids Research (2018) doi:10.1093/nar/gkx922.

50. Katoh, K. & Standley, D. M. MAFFT multiple sequence alignment software version 7: Improvements in performance and usability. Molecular Biology and Evolution 30, 772–780 (2013).

51. Capella-Gutiérrez, S., Silla-Martínez, J. M. & Gabaldón, T. trimAl: A tool for automated alignment trimming in large-scale phylogenetic analyses. Bioinformatics (2009) doi:10.1093/bioinformatics/btp348.

52. Darriba, D., Taboada, G. L., Doallo, R. & Posada, D. ProtTest-HPC: Fast selection of best-fit models of protein evolution. in Lecture Notes in Computer Science (including subseries Lecture Notes in Artificial Intelligence and Lecture Notes in Bioinformatics) vol. 6586 LNCS 177–184 (2011).

53. Kozlov, A. M. et al. RAxML-NG: A fast, scalable and user-friendly tool for maximum likelihood phylogenetic inference. Bioinformatics (2019) doi:10.1093/bioinformatics/btz305.

54. Minh, B. Q. et al. IQ-TREE 2: New Models and Efficient Methods for Phylogenetic Inference in the Genomic Era. Molecular Biology and Evolution (2020) doi:10.1093/molbev/msaa015.

55. Battye, T. G. G., Kontogiannis, L., Johnson, O., Powell, H. R. & Leslie, A. G. W. iMOSFLM: a new graphical interface for diffraction-image processing with MOSFLM. Acta Crystallogr D Biol Crystallogr 67, 271–281 (2011).

56. Evans, P. R. An introduction to data reduction: space-group determination, scaling and intensity statistics. Acta Crystallogr D Biol Crystallogr 67, 282–292 (2011).

57. Winn, M. D. et al. Overview of the CCP4 suite and current developments. Acta Crystallogr D Biol Crystallogr 67, 235–242 (2011).

58. French, S. & Wilson, K. On the treatment of negative intensity observations. Acta Crystallographica Section A 34, 517–525 (1978).

59. Potterton, L. et al. CCP4i2: the new graphical user interface to the CCP4 program suite. Acta Crystallogr D Struct Biol 74, 68–84 (2018).

60. McCoy, A. J. et al. Phaser crystallographic software. J Appl Crystallogr 40, 658–674 (2007).

61. Murshudov, G. N. et al. REFMAC5 for the refinement of macromolecular crystal structures. Acta Crystallogr D Biol Crystallogr 67, 355–367 (2011).

62. Emsley, P. & Cowtan, K. ıt Coot: model-building tools for molecular graphics. Acta Crystallographica Section D 60, 2126–2132 (2004).

63. Emsley, P., Lohkamp, B., Scott, W. G. & Cowtan, K. Features and development of Coot. Acta Crystallogr D Biol Crystallogr 66, 486–501 (2010).

64. Schüttelkopf, A. W. & van Aalten, D. M. F. ıt PRODRG: a tool for high-throughput crystallography of protein–ligand complexes. Acta Crystallographica Section D 60, 1355–1363 (2004).

65. Liebschner, D. et al. Macromolecular structure determination using X-rays, neutrons and electrons: recent developments in Phenix. Acta Crystallogr D Struct Biol 75, 861–877 (2019).

66. The PyMOL Molecular Graphics System, Version 2.4.1 Schrödinger, LLC.

67. Laskowski, R. A., MacArthur, M. W., Moss, D. S. & Thornton, J. M. ıt PROCHECK: a program to check the stereochemical quality of protein structures. Journal of Applied Crystallography 26, 283–291 (1993).

68. Davis, I. W. et al. MolProbity: all-atom contacts and structure validation for proteins and nucleic acids. Nucleic Acids Res 35, W375–W383 (2007).

69. Young, J. Y. et al. Worldwide Protein Data Bank biocuration supporting open access to high-quality 3D structural biology data. Database (Oxford) 2018, bay002 (2018).

70. Dong, R., Pan, S., Peng, Z., Zhang, Y. & Yang, J. mTM-align: a server for fast protein structure database search and multiple protein structure alignment. Nucleic Acids Res 46, W380–W386 (2018).

71. Saitou, N. & Nei, M. The neighbor-joining method: a new method for reconstructing phylogenetic trees. Molecular Biology and Evolution 4, 406–425 (1987).

72. Dolinsky, T. J., Nielsen, J. E., McCammon, J. A. & Baker, N. A. PDB2PQR: an automated pipeline for the setup of Poisson-Boltzmann electrostatics calculations. Nucleic Acids Res 32, W665–W667 (2004).

73. Tian, W., Chen, C., Lei, X., Zhao, J. & Liang, J. CASTp 3.0: computed atlas of surface topography of proteins. Nucleic Acids Res 46, W363–W367 (2018).

74. Laskowski, R. A., Jabłońska, J., Pravda, L., Vařeková, R. S. & Thornton, J. M. PDBsum: Structural summaries of PDB entries. Protein Sci 27, 129–134 (2018).

75. Bronner, V et al., Rapid and efficient determination of kinetic rate constants using the ProteOn XPR36 protein interaction array system, Bio-Rad bulletin 3172 (2006).

76. PyMol 2.1. https://github.com/schrodinger/pymol-open-source (2019).

77. Stewart, J. J. P. MOPAC2016-Stewart Computational Chemistry, Colorado Springs, CO, USA. 2016.

78. Stewart, J. J. P. Optimization of parameters for semiempirical methods VI: more modifications to the NDDO approximations and re-optimization of parameters. Journal of molecular modeling 19, 1–32 (2013).

79. Mato, J. & Guidez, E. B. Accuracy of the PM6 and PM7 Methods on Bare and Thiolate-Protected Gold Nanoclusters. The Journal of Physical Chemistry A 124, 2601–2615 (2020).

80. Unke, O. T., Koner, D., Patra, S., Käser, S. & Meuwly, M. High-dimensional potential energy surfaces for molecular simulations: from empiricism to machine learning. Machine Learning: Science and Technology 1, 013001 (2020).

81. Bushnell, E. A. C., Huang, W. & Gauld, J. W. Applications of Potential Energy Surfaces in the Study of Enzymatic Reactions. Advances in Physical Chemistry 2012, 867409 (2012).

