## Supplementary material for "Structural analysis unravels the functional promiscuity of Quinolone synthase-mediated polyketide biosynthesis in *Aegle marmelos* Correa": Supplementaryfigures.pdf

### Supplementary Figures

**Supplementary Figure S1: Sequence-structure alignment showing the conservation pattern of type III PKS from the Asgard species (*Candidatus Thorarchaeota archaeon*).** Alignment was prepared using ClustalW and ESPript 3.0 (MsCHS-CHS from *Medicago sativa*, AmQNS-Quinolone synthase from *Aegle marmelos*, AtCHS-CHS from *Arabidopsis thaliana*). The conservation level is shown by a color gradient (white-poor conservation, red-high conservation).

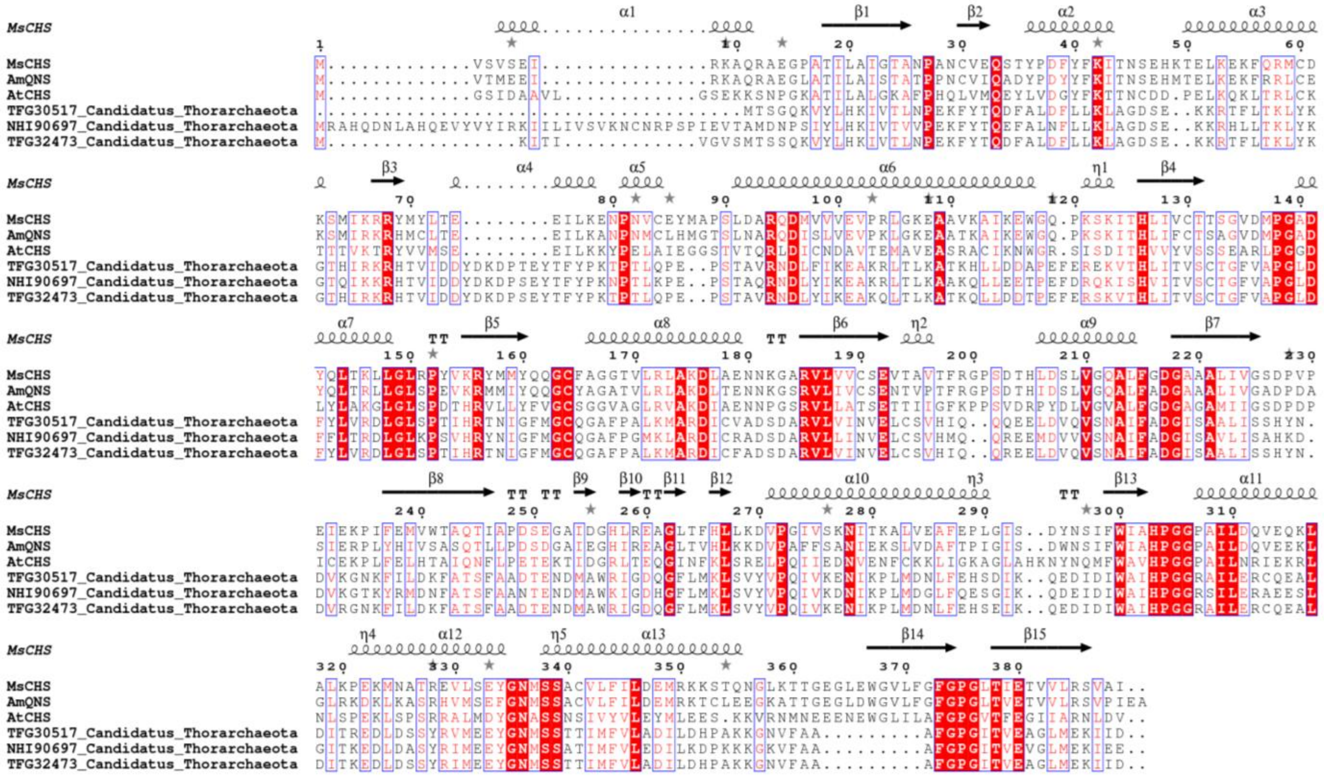

**Supplementary Figure S2. AmQNS - *N*-Methylanthraniloyl-CoA binding studies**.

AmQNS was successfully entrapped on the surface of a medium capacity 'GLM chip', and its interaction with its most favourable natural substrate, *N*-methylanthraniloyl-CoA (MANT-CoA), appears to fit well into the Langmuir model.

In the figure, **A)** L2 - reference channel. **B, C**) Channel L3 and L4 were used for AmQNS (50 µg/ml) immobilization (duplicate). SPR responses on the sensor chip surface (L3 - 9962 RU; L4 - ~7000 respectively, i.e., AmQNS was captured on the chip to the maximum final density of 9962 RU). **D, E** and **F**) analyte interaction (*N*-Methylanthraniloyl-CoA) to the immobilized AmQNS at various concentrations and its kinetics. Not all concentrations are shown. **G)** Association (ka), dissociation (kd), and equilibrium (K) constants derived from the SPR sensogram for AmQNS- different CoA interactions

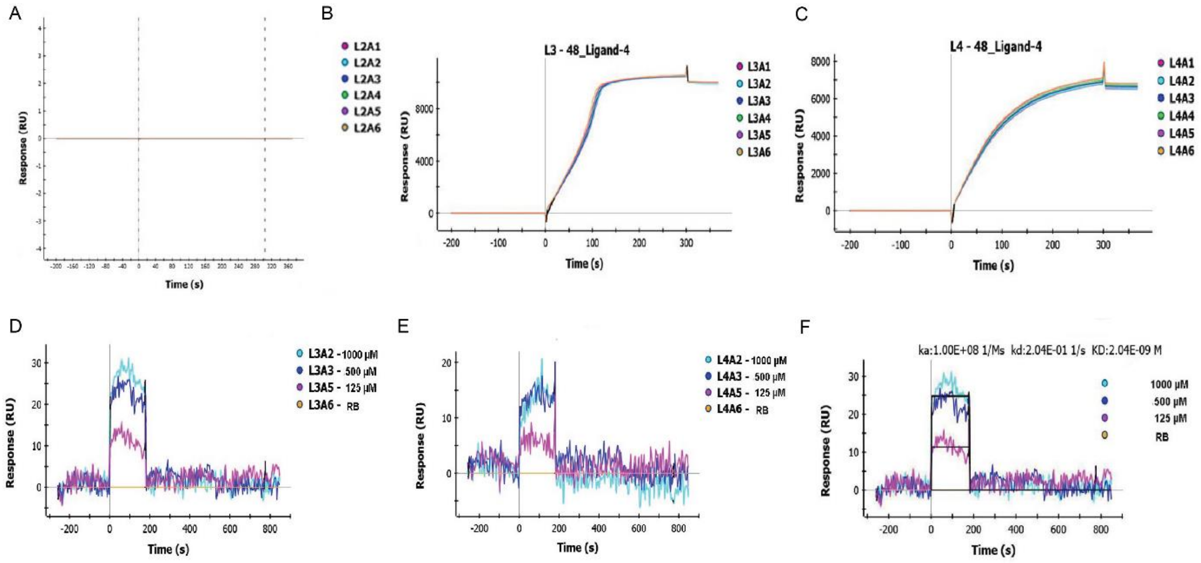


G

| No | Substrate | ka (1/Ms) | kd (1/s) | K D (M) |
|---|---|---|---|---|
| 1 | N-Methylanthraniloyl-CoA | 1.00E+08 | 2.04E-01 | 2.04E-09 |
| 2 | P-Coumaroyl-CoA | 5.91E+05 | 5.95E-02 | 1.01E-03 |
| 3 | Feruloyl-CoA | 1.00E+08 | 9.83E-01 | 9.83E-09 |
| 4 | Hexanoyl-CoA | 9.63E+05 | 7.03E-03 | 7.30E-09 |
| 5 | Palmitoyl-CoA | 4.58E+04 | 4.17E-03 | 9.11E-08 |
| 6 | Benzoyl-CoA | 9.51E+03 | 2.67E-02 | 2.80E-06 |
| 7 | Myristoyl-CoA | 1.30E+04 | 4.04E-03 | 3.10E-07 |
| 8 | Butyryl-CoA | 3.61E+04 | 4.95E-02 | 1.37E-06 |
| 9 | Stearoyl-CoA | 1.17E+04 | 1.93E-03 | 1.64E-07 |
| 10 | Malonyl CoA | 7.82E+04 | 1.34E-02 | 1.72E-07 |
| 11 | Sinapoyl-CoA | 1.41E+05 | 1.63E-02 | 1.16E-07 |

**Supplementary Figure S3**. Structure summary of AMQNS. Modified PDBsum output showing the secondary structure wiring diagram.

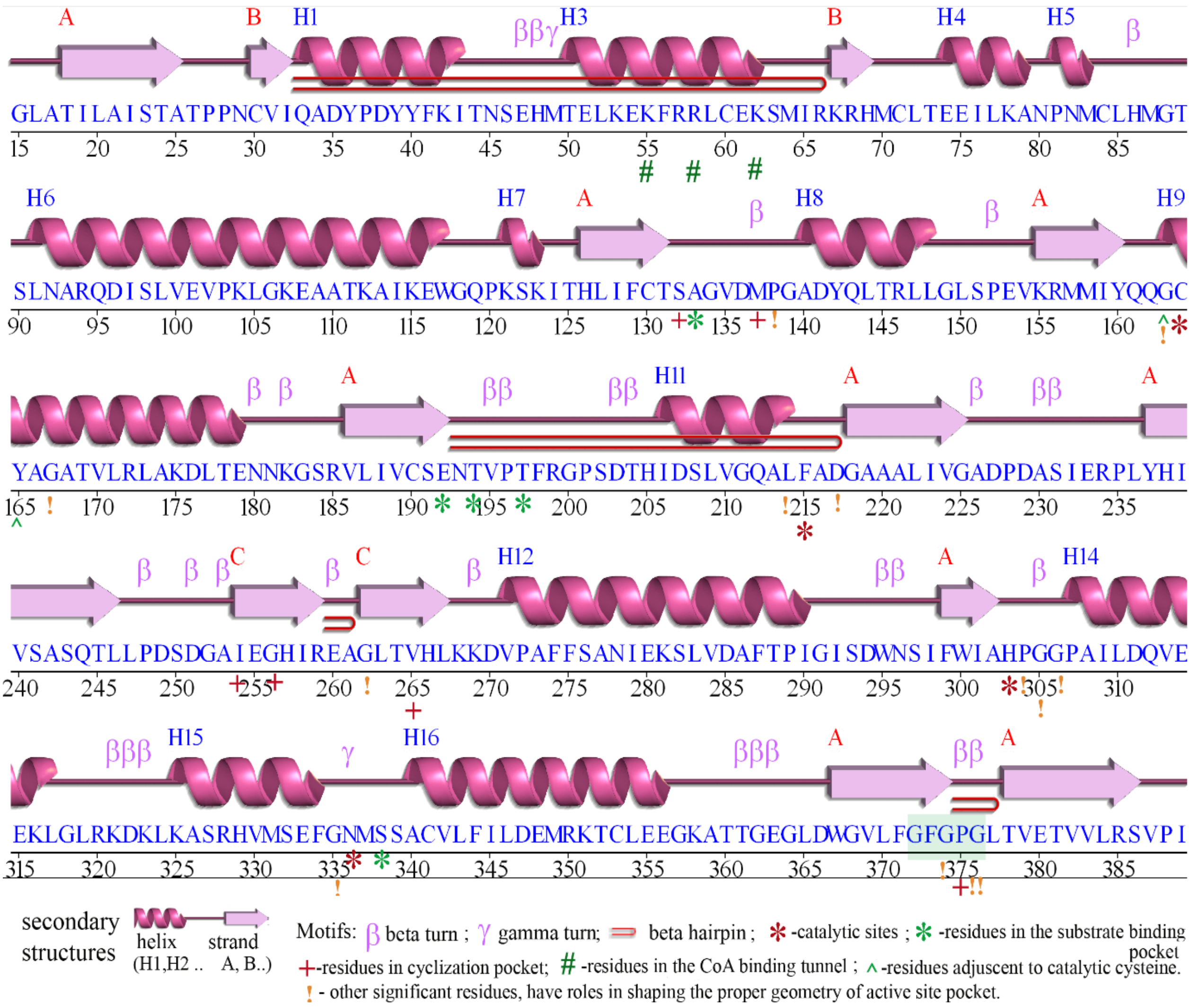

**Supplementary Figure S4.** A) Comparison of cavity parameters of AmQNS with nearest homologues B) Cartoon representation of AmQNS, CmACS (3WD7) and CmQNS (3WD8)

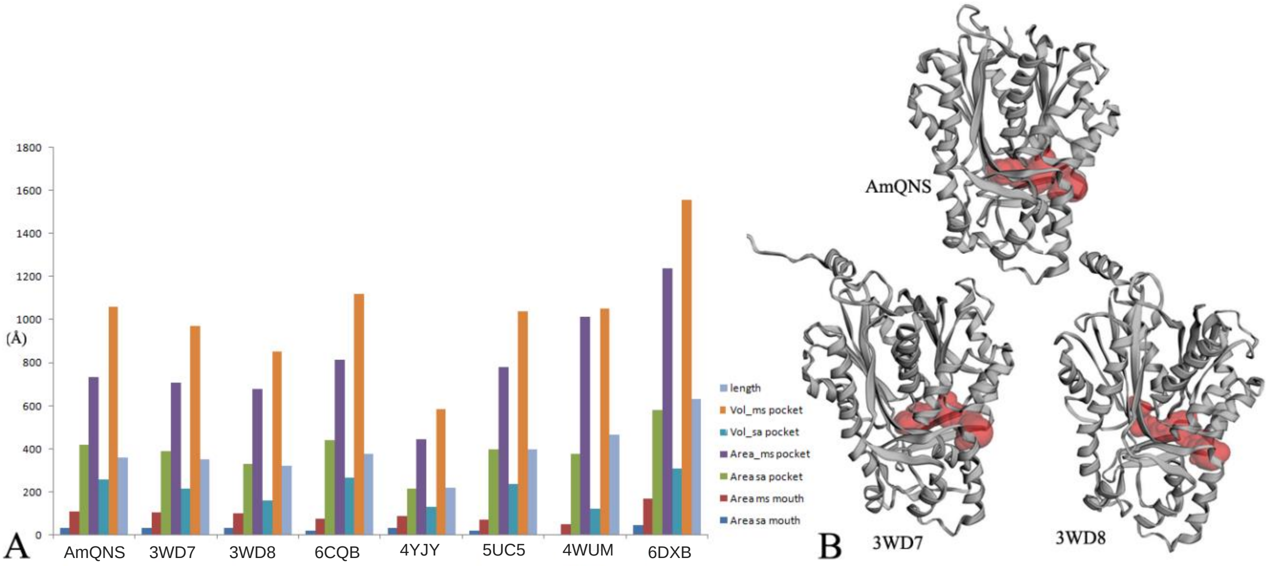


**Supplementary figure S5**

Cartoon representation of active site architecture of AmQNS and nearest homologues.

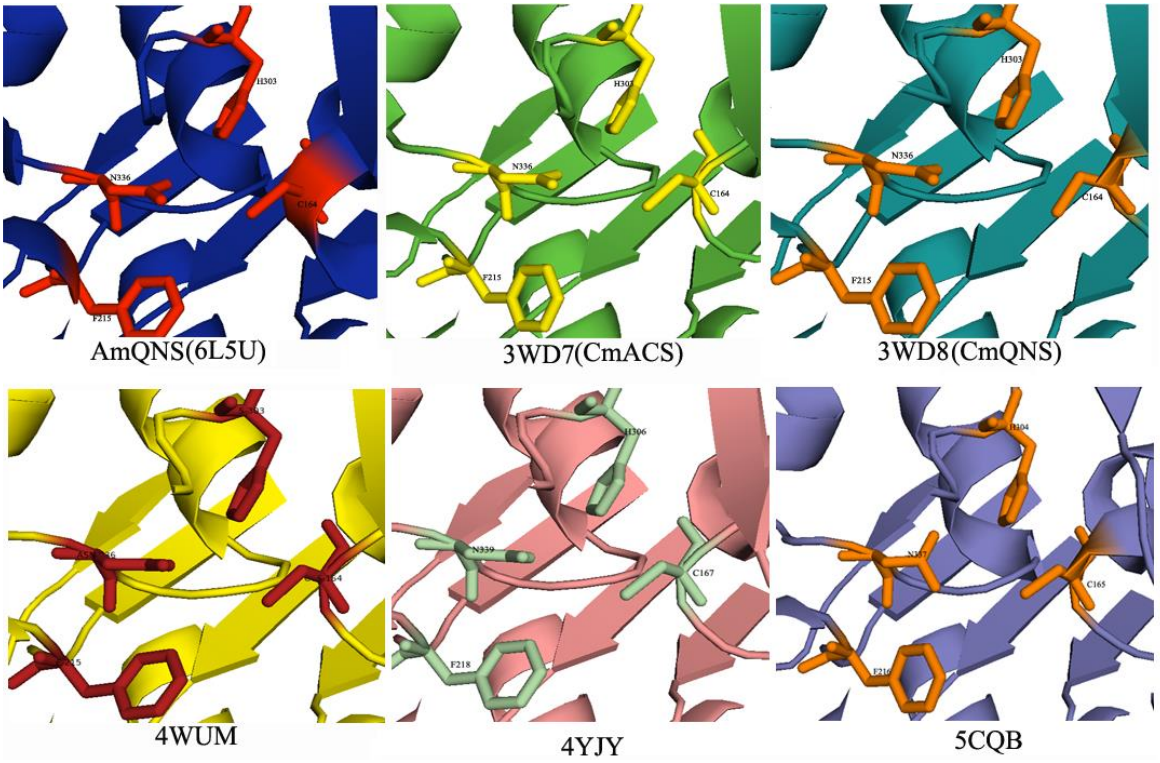

**Supplementary figure S6.** 2Fo-Fc electron density map of the active site residues and substrate binding residues in the superimposed models (contoured at 1.0 sigma level). A) AmQNS superimposed with CmACS. Active site residues of AmQNS represented in orange color, and of CmACS in pink color B) Substrate binding residues in the superimposed model C) superimposition of AmQNS with CmACS and CmQNS.

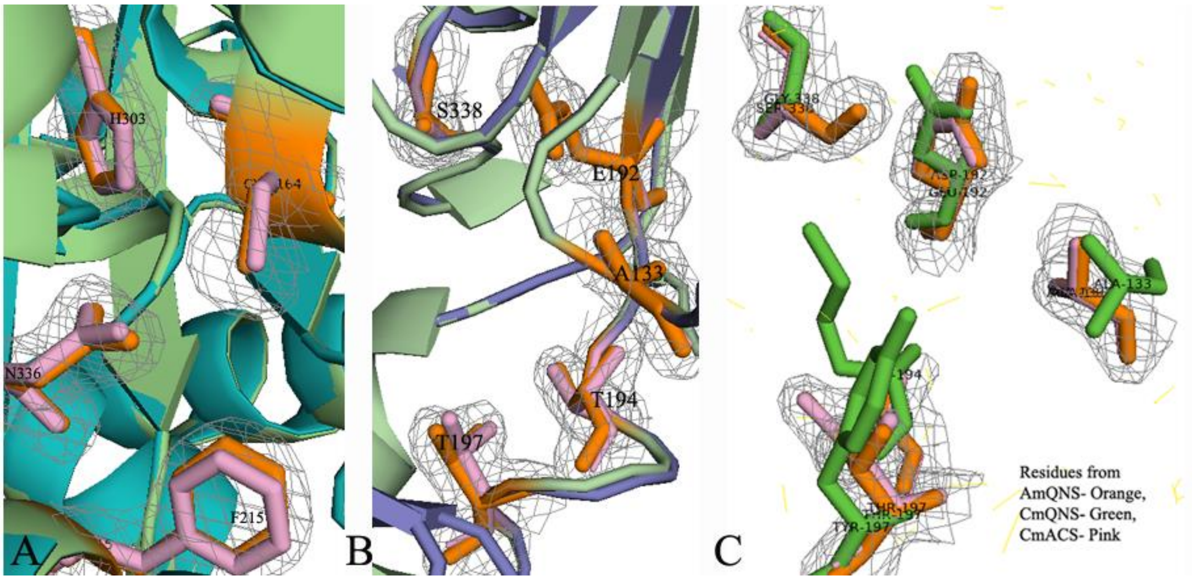

**Supplementary Figure S7:** CoASH (by-product)-bound AmQNS. Orientation of the ligand molecule in the cavity is depicted. Significant residues highlighted (catalytic sites (C-F-H-N ) in orange; substrate binding sites in pink, cyclization residues in yellow, CoA binding residues in pale cyan, Residues shaping the geometry of active sites in blue. B) Enlarged view of substrate binding pocket above and the position and orientation of significant residues, upon ligand binding is below C) Ligplot representation of AMQNS - CoASH interactions. Hydrogen bonds between protein and ligands are highlighted in green dashed lines.

A

B

C

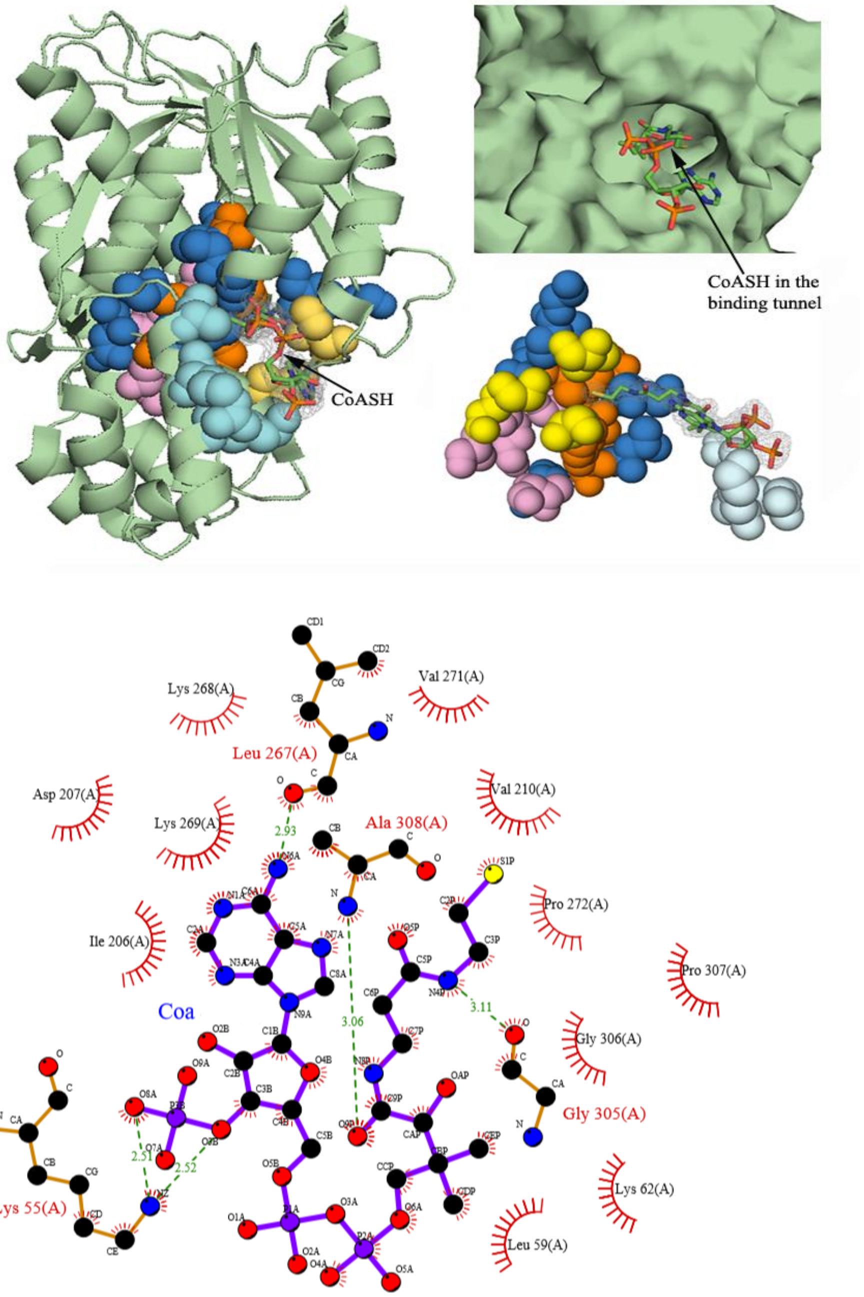

**Supplementary Figure S8:** Chemical structure of MANT-CoA and derivatives, quinolone and acridones.

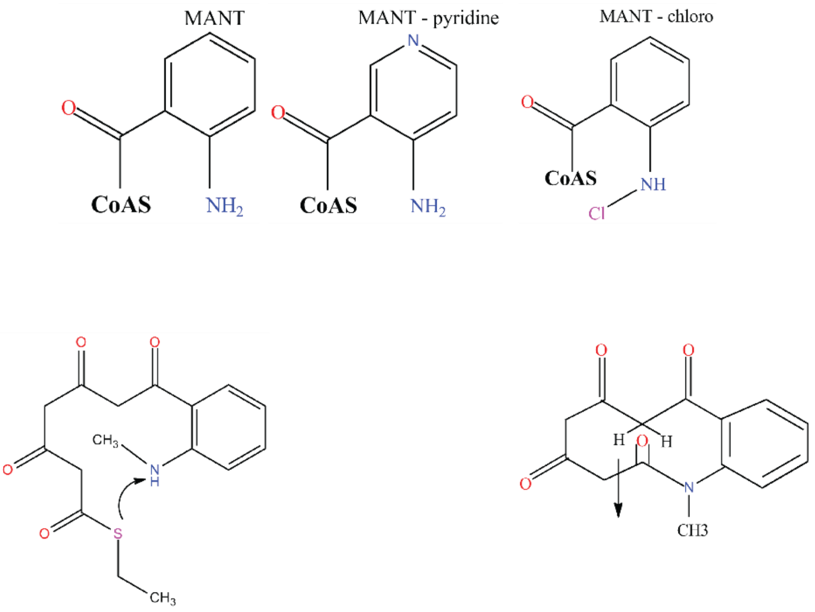
